# Local IL-17 orchestrates skin aging

**DOI:** 10.1101/2022.01.31.478459

**Authors:** Paloma Solá, Elisabetta Mereu, Júlia Bonjoch, Marta Casado, Oscar Reina, Enrique Blanco, Manel Esteller, Luciano DiCroce, Holger Heyn, Guiomar Solanas, Salvador Aznar Benitah

## Abstract

Skin aging is characterized by structural and functional changes that lead to slower wound healing and higher rate of infections, which contribute to age-associated frailty. This likely depends on synergy between alterations in the local microenvironment and stem cell–intrinsic changes, underscored by pro-inflammatory microenvironments that drive pleotropic changes. To date, little is known about the precise nature and origin of the proposed age-associated inflammatory cues, or how they affect different tissue resident cell types. Based on deep single-cell RNA-sequencing of the entire dermal compartment, we now provide a comprehensive understanding of the age-associated changes in all skin cell types. We show a previously unreported skew towards an IL-17–expressing phenotype of Th cells, γδ T cells and innate lymphoid cells in aged skin. Aberrant IL-17 signaling is common to many autoimmune (e.g., rheumatoid arthritis and psoriasis) and chronic inflammatory diseases. Importantly, *in vivo* blockade of IL-17–triggered signaling during the aging process reduces the pro-inflammatory state by affecting immune and non-immune skin cells of both dermis and epidermis. Strikingly, IL-17 neutralization significantly delays the appearance of age-related traits, such as decreased epidermal thickness, increased cornified layer thickness and ameliorated hair follicle stem cell activation and hair shaft regeneration. Our results indicate that the aged skin shows chronic and persistent signs of inflammation, and that age-associated increased IL-17 signaling could be targeted as a strategy to prevent age-associated skin ailments in elderly.

## Introduction

Aging is characterized by an accumulation of cellular damage due to an imbalance between damage generation and clearance, in part due to an ineffective metabolism, circadian clock rewiring, increased systemic inflammation and accumulation of senescent cells^1–6^. This creates the perfect scenario for malfunctioning of the organism and eventual collapse when facing a challenge.

The skin contains a multilayered epidermis interspersed with mini-organs (i.e. hair follicles [HFs] and sebaceous glands) embedded in it. The outermost layer of skin, the epidermis, is a stratified epithelium that forms an impermeable protection for the organism. Daily renewal of the epidermis is fueled by interfollicular epidermal (IFE) stem cells (IFE-SCs), whereas hair follicle-stem cells (HF-SCs) maintain the hair follicles by periodically generating new hair shafts^7^. In the dermis, an organized mesh of extracellular matrix (ECM) embedded with several types of cell lineages form the niche for all types of keratinocyte stem cells^7^. Aging causes architectural and functional changes in the epidermis that affect its regenerative potential and its function as a barrier^8–12^. Moreover, age-related changes in the cellular composition and ECM properties of the dermis, which are still ill-defined, affect the function of the dermis as a structural scaffold and as a niche for epidermal stem cells^13–18^. Collectively, these changes lead to a slower epidermal turnover, more breaching of the barrier, and a lower quality wound healing, all of which contribute to increasing the incidence of infections and chronic wounds in the elderly^19–21^. In sum, there are dual characteristics of aging on skin homeostasis: i) it affects the interfollicular epidermal stem cells and hair follicle stem cells in a cell-autonomous manner; and ii) it leads to misregulation of the delicate dialogue between these stem cells and their subjacent niches in the dermis.

Single cell RNA-seq efforts have been recently directed towards defining the relationship between the different skin cell types during homeostasis, aging and disease ^13,22–27^ These studies suggest that some immune cell types change their abundance or behavior during aging in skin, including regulatory T cells (T-regs), dendritic epidermal T cells and Langerhans cells^28–31^. Such changes are especially relevant, considering that the dialogue between immune cells and the epidermis is essential for wound healing ^28,32^. However, much is still unknown regarding the relationship between immune cells and non-immune cells of the dermis and the epidermis during aging.

Here, we have unbiasedly profiled and characterized to an unprecedented depth the single-cell transcriptome of dermal cells in aged skin in mice. Many cell types showed significant changes in their proportion and gene expression profile during aging; however, we functionally characterized dermal CD4+ T helpers, γδ T cells and innate lymphoid cells (ILCs), as those showing the most prominent transcriptional changes during aging that appeared to orchestrate many of the alterations observed in several other skin cell types. Specifically, these cell types showed a polarization towards an IL-17– producing phenotype, strongly contributing to the inflammatory environment found in aged skin. Importantly, in vivo blockade of IL-17 signaling during the aging process delayed the development of several hallmarks of skin aging, such as an amelioration of hair follicle regeneration, lack of thinning of the epidermal layer and a decreased cornified layer thickness, as well as a reduced epidermal inflammatory state.

## Results

### Aged dermal cells reveal cell type–specific, age-related changes

We analyzed the non-epithelial (e.g., negative for epithelial cell adhesion molecule [EpCAM–]) dermal population of dorsal skin from aged (80- to 90-week-old) and adult (17- to 25-week-old) mice, by single-cell RNA-sequencing (scRNA-seq). We first removed epidermal cells enzymatically and then we isolated dermal cells by fluorescence-activated cell sorting (FACS) (see workflow, Fig. 1a and Supplementary Fig. 1a). In order to maximize sampling of the less abundant immune cells, we enriched by FACS for CD45+ cells and sequenced them separately from the rest of dermal cells (EpCAM–/CD45–) (Supplementary Fig. 1a). Using the 10× Genomics platform (version 3), we characterized 11,940 cells for the CD45+ cells and 5,213 for their CD45– /EpCAM– counterparts. After batch correction and data integration, we clustered all cells together by generating a shared nearest neighbor graph using the Louvain algorithm^33^. To visualize the clustering, we used Uniform Manifold Approximation and Projection (UMAP), which verified good sample mixing. We performed differential gene expression between cell populations to obtain cluster markers (Fig. 1b and Supplementary Table 1) and plotted discriminatory population markers to ensure correct clustering (Fig. 1c).

**Figure 1.**
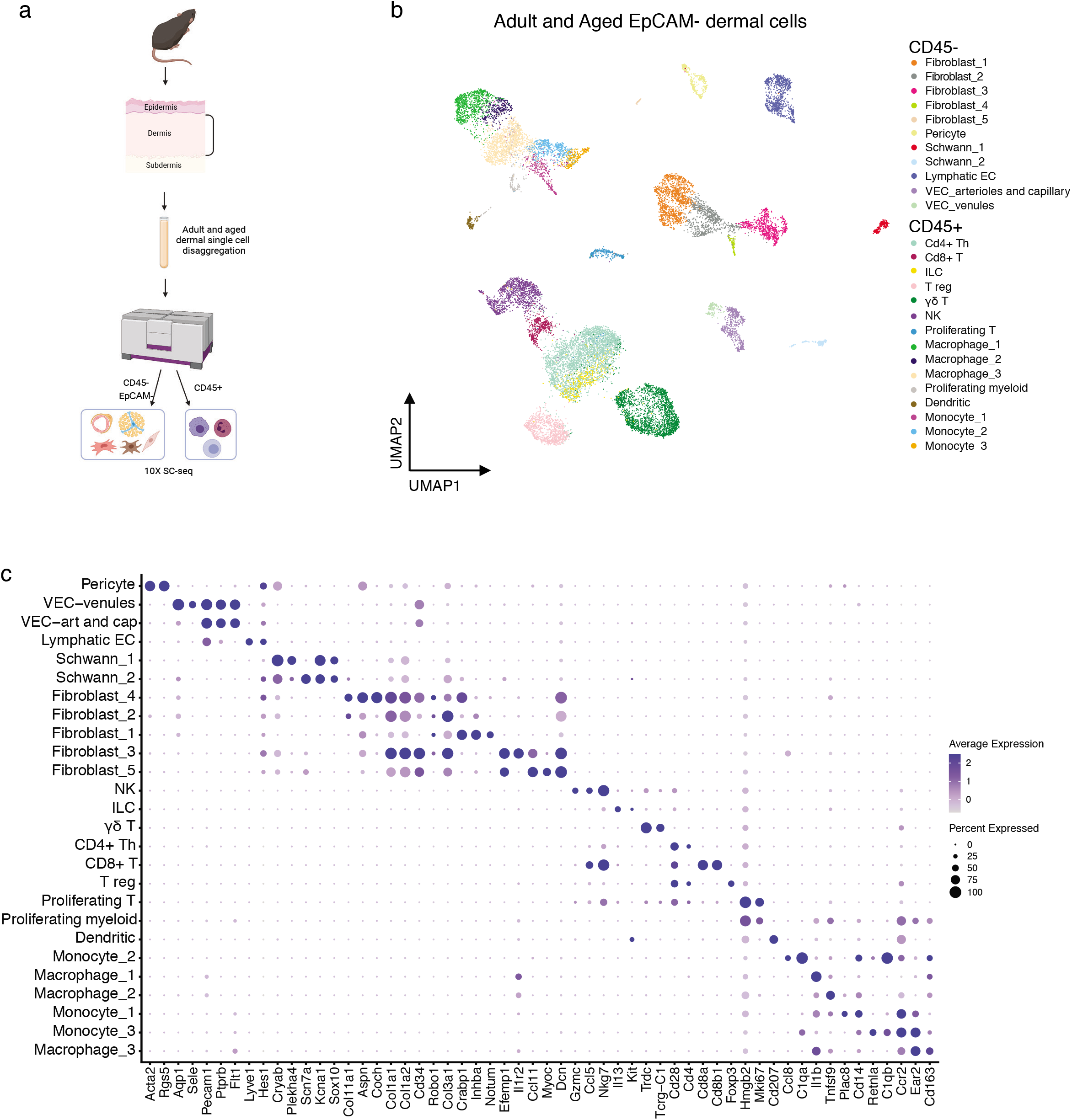
Dermal cell characterization by scRNA-seq. **a,** Workflow used to obtain dermal cells of adult and aged murine backskin. The single-cell suspensions were enriched for EpCAM–/CD45– and CD45+ cells separately by fluorescence activated cell sorting (FACS). The transcriptomes of sorted single cells were then analyzed by 10× scRNA-seq. For the CD45+ cells, *n* = 6 mice for the adult group, and *n* = 4 mice for the aged group, with three technical replicates; for the CD45–/EpCAM– cells, *n* = 2 mice for the control group, and *n* = 2 mice for the aged group, with two technical replicates. **b**, UMAP visualization of all adult and aged dermal cells analyzed by single-cell RNA-sequencing. **c**, Dot plot showing discriminatory markers for each cell type, subtype or state found in panel b.

The dermis is composed of several cell types that serve as structural and functional support for the skin^7^. In the UMAP visualization, we observed three major groups of clusters pertaining to non-immune (CD45–/EpCAM–), immune lymphoid and immune myeloid lineages (both CD45+) (Fig. 1b). Fibroblasts were the most abundant CD45– /EpCAM– cell type and separated in five clusters that contained subtypes with distinct features (Fig. 2a). Some of these clusters showed high similarities to well-defined fibroblast subtypes in terms of marker expression: cluster 2 was papillary-like; cluster 1, reticular-like; and cluster 3, pro-inflammatory^23,24,34,35^. These clusters differentially expressed distinct collagen types, as well as other ECM proteins, conferring them with distinct properties necessary to generate and maintain the dermal niche. Importantly, cluster fibroblast_1 expressed markers of dermal papilla cells^36^, a specialized cluster of fibroblasts located immediately beneath hair follicles that is essential for modulating the hair follicle cycle (Figs. 1c and 2a). We also detected clusters for lymphatic and vascular endothelial cells (ECs), along with pericytes and Schwann cells, in the CD45–/EpCAM– population (Figs. 1b,c and 2a). On the other hand, the CD45+ clusters of immune cells separated into two large groups based on their lymphoid or myeloid lineage, with several cell types and states among them (Fig. 1b,c).

**Figure 2.**
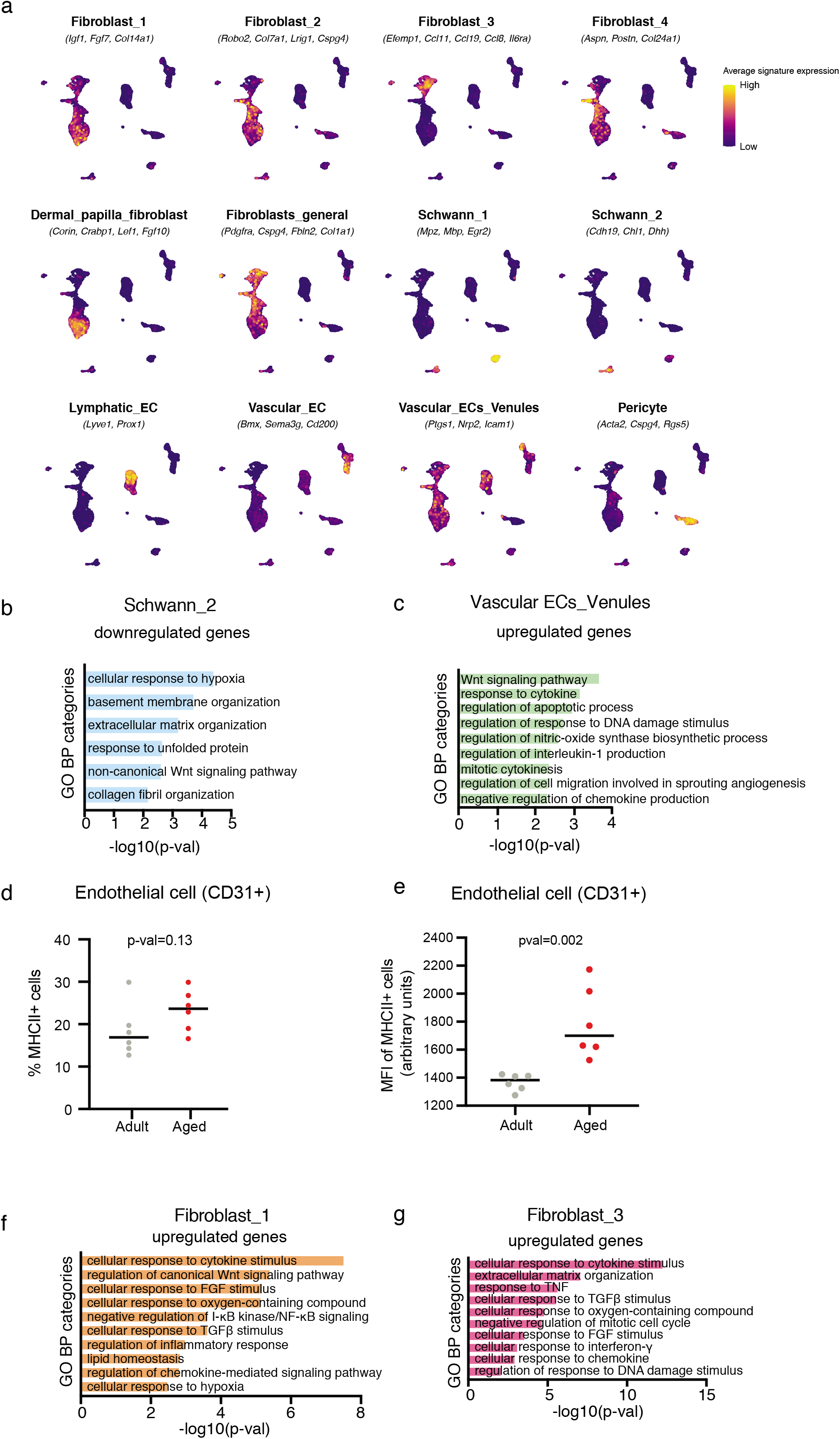
Aged non-immune cells show a shift towards age-associated and pro-inflammatory functions. **a**, UMAP visualization of cell subtype-specific signatures. **b**, Plot of selected gene ontology (GO) categories belonging to biological processes (BP) analysis for genes downregulated during aging in cells included in Schwann_2 cluster. The x axis represents –(log_10_) of the *P*-value for each depicted GO category. **c**, Plot of selected GO categories belonging to BP analysis for genes upregulated during aging in venule endothelial cells. The x axis represents –(log_10_) of the *P*-value for each depicted GO category. **d**, Percentage of endothelial cells (labeled by CD31+ expression) with MHCII expression on their surface detected by staining with fluorescently labeled antibodies and analyzed by flow cytometry. **e**, Mean fluorescence intensity (MFI) of MHCII staining in CD31+/MHCII+ dermal cells. For **d** and **e**, each data point shows individual value per mouse (*n* = 6 mice per age group); bars indicate median. *P*-values were obtained with Mann Whitney *U* test. **g**,**h**, Plot of selected GO categories belonging to BP analysis for genes upregulated during aging in fibroblasts included in clusters fibroblast_1 (**f**) and fibroblast_3 (**g**). The x axis represents –(log_10_) of the *P*-value for each depicted GO category.

To gain an overview of the transcriptomic changes induced by aging, we compared the 100 top marker expression in the clusters between adult and aged cells and plotted the Jaccard index between the adult and aged cells in each cluster. This analysis indicated that different cell types showed a progressive range of age-associated changes, from very small ones (e.g. monocyte_3 and lymphatic ECs) to major changes (e.g. ILCs or CD8+ T cells) (Supplementary Fig. 1b). Strikingly, this indicated that the transcriptional changes accumulated during aging were cell type–specific and varied between cell lineages even within the same tissue. Due to this, we focused on the age-associated changes separately for the different groups of cell clusters and the cell states obtained in our analysis.

### Distinct non-immune cells in aged skin show changes related to defective functions and increased inflammation

#### Schwann cells

We identified two clusters of Schwann cells within the CD45–/EpCAM– fraction (Fig. 1c and 2a, and Supplementary Table 1). Schwann cells are neural crest–derived glial cells of the peripheral nervous system^37^ that are important for regenerating injured axons as well as for initiating and promoting the healing of damaged peripheral tissues (including skin)^38–40^. Schwann cells need to undergo epithelial-to-mesenchymal transition (EMT) and migrate into the wounded area to exert their correct regenerative function. There, they remodel the ECM around the damaged nerves and proceed to repair them^41,42^. Their primary location in skin is at the dermal–epidermal junction, where they serve as initiators of pain and thermal sensation^43^. Importantly, aged mice exhibit reduced regeneration of peripheral nerves, such as the sciatic nerve, partially due to decreased Schwann cell functioning^44,45^. Little is known, however, about how the specialized cutaneous Schwann cells are affected during aging at the molecular level. The two clusters of Schwann cells that we observed represented a myelinating-differentiated (Schwann_1) *versus* a more precursor-like state (Schwann_2) (Fig. 2a). Interestingly, genes relevant for EMT, ECM remodeling and myelination (such as *Ezh2, Itga4, Nf1*) along with several types of collagen genes became downregulated in Schwann_2 of aged dermis (Supplementary Table 2), which was confirmed by gene ontology (GO) analysis (Fig. 2b and Supplementary Table 3). Altogether, our data suggested that aged Schwann cells could be less efficient in their regenerative functions in skin (Fig. 2b and Supplementary Table 3).

#### Endothelial cells

We detected three clusters of endothelial cells (ECs): one corresponding to lymphatic ECs, and two representing vascular ECs (arteriole and capillary ECs in one cluster, and venule ECs in the other) (Fig. 1c and 2a). Microcirculation constitutes the blood flow through the network of small tissue-embedded microvessels. During aging, microvessels —arterioles, capillaries and venules— show decreased density and higher stiffness, accompanied by decreased reactivity to stimuli. These changes in blood flow negatively impact skin oxygenation and nutrient delivery, and lead to deregulation of the inflammatory responses, among other consequences^46^. Like all blood and lymph vessels, microvessels are lined by endothelial cells (ECs). Such vascular ECs constitute key players in the immune response and are among the first line of cells that detect external pathogens in the bloodstream and become activated by them. Specifically, activated venules secrete cytokines and chemokines to recruit immune cells and even have a “nonprofessional” antigen-presenting function during inflammation^47^; in chronic inflammation, activated venules can upregulate the expression of the class II major histocompatibility family of proteins (MHCII), one of the key molecules for antigen presentation to T cells^48^. Interestingly, we found that aged venule ECs showed higher expression of genes involved in cytokine production and signaling, pointing to a more inflammatory state associated to skin aging (Fig. 2c and Supplementary Table 2 and 3). These gene expression changes were reminiscent to those associated to diseased states, such as psoriasis^25^. We then analyzed levels of MHCII proteins in the surface of adult and aged ECs by flow cytometry. Of note, although the number of CD31+ cells with MHCII on their surface increased with age, this was still not significant (Fig. 2d). However, the levels of MHCII on the surface of these CD31+/MHCII+ cells were significantly elevated in aged tissue (Fig. 2e). Taken together, our results indicate that venule ECs undergo a shift towards a more pro-inflammatory state, with features of non-professional antigen– presenting cells, reminiscent of autoimmune diseases. This observation is in line with previous findings regarding the higher pro-inflammatory state of aged ECs in the cardiovascular system^49^.

#### Fibroblasts

During aging, mouse dermal fibroblasts reduce their production of ECM components, increase their expression of inflammation-related genes and acquire traits of adipocytes^5,18,22,34^. We did not observe notable changes in the proportions of any of the five fibroblast subpopulations in the aged skin as compared to the adult tissue (Supplementary Fig. 2a). However, cluster fibroblast_1 (corresponding to reticular-like fibroblasts) showed an upregulation of genes related to cytokine, chemokine and inflammatory responses during aging (Fig. 2f and Supplementary Table 3). Alterations in expression of pro-inflammatory genes were even more pronounced in fibroblasts in the cluster fibroblast_3, which define fibroblasts involved in inflammatory responses in the adult skin (Fig. 2a,g and Supplementary Fig. 2b).

### Increased IL-17–related inflammation in the immune compartment of aged skin

Altogether, our results indicated an overall increase of inflammation as the predominant features in non-immune cells in the aged skin. To further dissect the origin of these changes, we subsequently focused on the immune cells, which are a major source of pro-inflammatory cytokines. We identified two major groups of clusters in our analysis based on lineage of either lymphoid or myeloid (Fig. 3a).

**Figure 3.**
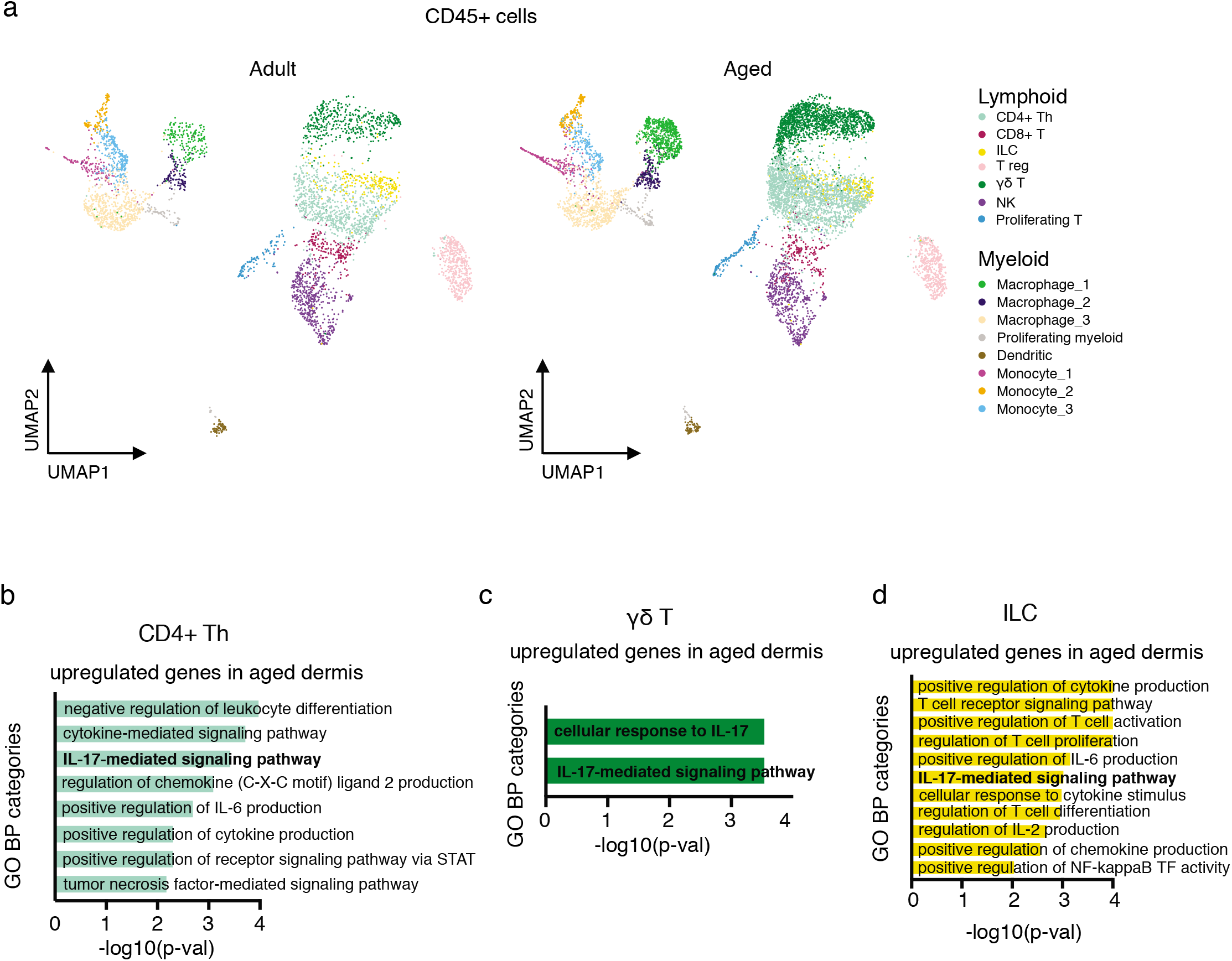
Lymphoid immune cells show increased IL-17A/F-related signaling. **a**, UMAP representation of immune (CD45+) cells. Left panel, adult cells, and right panel, aged cells. **b**–**d**, Plot of selected GO categories belonging to BP analysis for genes upregulated during aging in CD4+ Th cells (**b**), γδ T cells (**c**) and innate lymphoid cells (ILC) (**d**). The x axis represents –(log_10_) of the *P*-value for each depicted GO category. Highlighted in bold are the IL-17-related GO categories.

#### Dermal myeloid cells

Dermal myeloid cells balance pro- and anti-inflammatory functions in skin in a contextdependent manner^50^. Specifically, they scan the tissue to detect antigens, orchestrate an early response to pathogens and trigger the first steps in the wound healing response^51^. We detected eight clusters of myeloid cells containing different subtypes of macrophages (macrophage_1 to _3), monocytes (monocyte_1 to _3), dendritic cells (DCs), and proliferating myeloid cells (Fig. 3a).

Monocyte_1 and macrophage_1 clusters were defined by pro-inflammatory features (e.g., among the markers for monocyte_1 we found *Ccr2, Cxcl2, Fcgr4, Cxcr4* for; *Il1b* and *Tnfaip3;* and for macrophage_1, MHCII complex genes such as *H2Ab1, H2-Eb1. H2-Aa* and *CD74*). Conversely, cluster macrophage_2 expressed markers of regulatory macrophages (such as *Il4l1, Cd200r1* and *Lgals1*) (Supplementary Table 1). Of note, although the proportion of both macrophage_1 and macrophage_2 clusters increased in the aged tissue (Supplementary Fig. 2a), they showed no clear changes in their transcriptomes, indicating that aging induced alterations in their numbers rather than their state (Supplementary Table 2). Conversely, monocyte_1 and monocyte_2 clusters showed more pronounced changes in gene expression with aging, with a clear trend towards expressing higher levels of pro-inflammatory cytokine secretion and responses to pro-inflammatory stimuli (Supplementary Fig. 2c and Supplementary Tables 2 and 3); this included genes encoding IL1β and TNF, which are necessary for these immune cells to initiate a full immune response^51^. Interestingly, the expression of these genes also was upregulated in clusters macrophage_2, monocyte_1, monocyte_2 and dendritic cells during aging (Supplementary Table 2). Altogether, this indicates that myeloid cells exacerbate a chronic inflammatory state in aged skin.

#### T cells and ILCs

Lymphoid cells in the mouse dermis include several types of T cells (CD4+, CD8+, γδ T cells and regulatory T cells [T-regs]), innate lymphoid cells (ILCs) and natural killer cells (NKs), which vary in their abundance in homeostatic conditions (Fig. 3a, left panel). These cells scan for tissue damage and perform immune surveillance in steady-state, as well as trigger an inflammatory response during wound healing and tumorigenesis^50^. Strikingly, the proportion of γδ T cells and CD4+ T helper cells increased in aged dermis (Fig. 3a, right panel, and Supplementary Fig. 2a). Moreover, these cells types were among those with a higher proportion of cells affected by aging (Supplementary Fig. 2d).

Furthermore, in the aged tissue, γδ T cells, CD4+ T helper cells and ILCs expressed much higher levels of inflammatory genes when compared to their adult counterparts; interestingly, these genes were predominantly related to the pro-inflammatory cytokine interleukin 17 (IL-17) (Fig. 3b,c,d and Supplementary Tables 2 and 3).

There are six IL-17 members, IL-17A to -F; all of these signal via binding to the IL-17R family of receptors. Of these, IL-17A and IL-17F are highly homologous and can function as heterodimers^52^. The IL-17 family of cytokines performs a plethora of activities, ranging from tissue repair and host defense, to pathogenic ones, such as in autoimmune and chronic inflammatory diseases^52^. Importantly, therapeutic inhibition of aberrant IL-17A and IL-17F activities is used as treatment for skin diseases, such as psoriasis, pemphigus and other autoimmune conditions^53–56^. Interestingly, during aging there is a polarization of circulating CD4+ T helper cells towards an IL-17 expressing phenotype, and in mice, γδT cells also show this skew in peripheral lymph nodes ^57–59^. However, whether these changes affect peripheral tissue aging (and if so, how they do it) is unknown.

As mentioned above, we observed an increased expression of *Il17a* and *Il17f* in individual CD4+ Th cells, γδ T cells and ILCs (Fig. 3b,c,d and Supplementary Table 2), suggesting that there was a polarization towards an IL-17–expressing phenotype (Fig. 4a). Zooming in the CD4+ Th cluster further revealed three subclusters with specific marker gene expression (Fig. 4b,c, and Supplementary Table 1). While cluster CD4+ Th_a could not be assigned to any specific Th subtype, the CD4+ Th_c cluster showed markers usually expressed by T_H_1 cells, such as *Ifng*. Interestingly, the cluster CD4+ Th_b showed markers compatible with being *bona fide* T_H_17 cells, such as *Il17f, Rora, Tmem176a/b, Ccr6* and *JunB*, among others^60–62^ (Fig. 4c). This cell subtype, CD4+ Th_b (T_H_17), constituted the most remarkably abundant in aged dermis compared to control adult skin (Fig. 4d).

**Figure 4.**
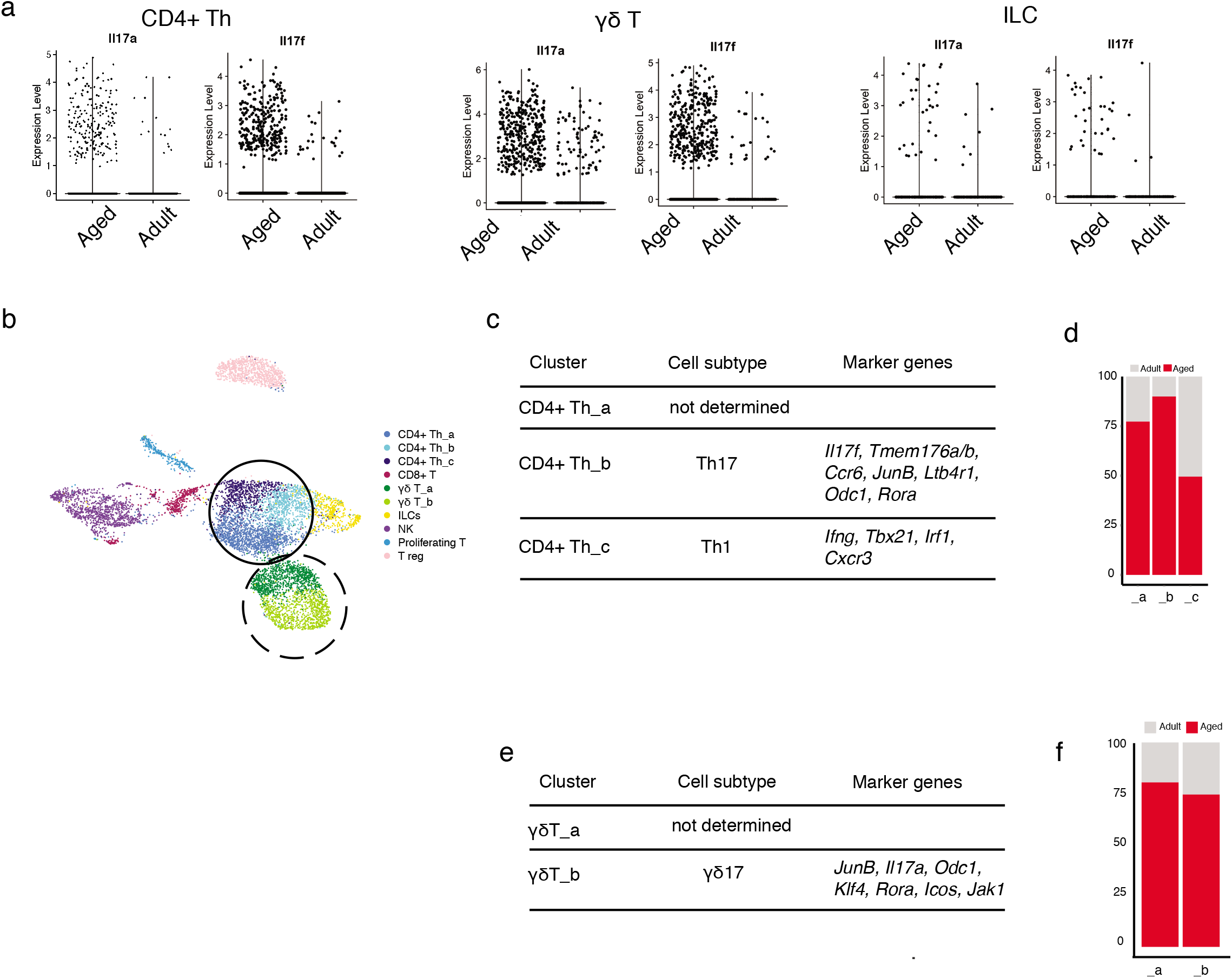
Lymphoid skew towards an IL-17 expressing phenotype upon aging. **a**, Violin plots comparing the expression values of *Il17a* and *Il17f* in aged and adult CD4+ Th, γδ T cells and innate lymphoid cells (ILC). **b**, UMAP representation of subclustering of the previously described CD4+ Th and γδ T clusters. Three new clusters of CD4+ Th were found (CD4+Th_a, _b and _c), shown here in blue and circled by the continuous line; and two new clusters of γδ T were found (γδ T_a, and _b), in green and circled by the dashed line. **c**, List of relevant markers that discriminate the subclusters belonging to specific CD4+ Th subtypes. **d**, Bar plot with the proportions of these subclusters in aged and adult samples. **e**, List of relevant markers that discriminate the subclusters belonging to specific γδ T cell subtypes. **f**, Bar plot with the proportions of the γδ T cell subclusters in aged and control groups.

Similar to Th cells, γδ T cells could also be further clustered into two subpopulations whose proportion increased in aged skin (Fig. 4c,f and Supplementary Table 1). Cluster γδT_b, expressed markers compatible with γδ T17 cells as indicated by the expression of *Il17a, Rora, JunB* and *Jak1*, among others^62^, whereas γδT_a cluster could not be assigned to any specific γδ T cell subtype (Fig. 4e). Both γδ T clusters were more abundant in aged skin, showing an increased presence of γδ T cells and more importantly IL-17-expressing γδ T cells in the aged murine skin.

ILCs are crucial for the development of psoriatic pathogenesis through sustained increased secretion of IL-17^63^. Our analysis also showed a marked increase in the expression of genes associated to the switch from ILC subtype to ILC3^63^, such as *Il17a*, *Il17f*, *Rora* and *Ccr6*, in aged skin (Fig. 4a and Supplementary Table 2).

### Inhibition of IL-17A/F signaling prevents age-related inflammation in dermal cells

We next sought to study whether the significant increase in the expression of *Il17a* and *Il17f* in a subset of aged lymphoid cells contributes to the general pro-inflammatory state of the aged skin. Treatment with anti-IL-17A/F antibodies is currently used as a therapy for patients with inflammatory and autoimmune diseases, such a psoriasis^56^. We therefore blocked IL-17 signaling by systemically administering neutralizing antibodies against IL-17A and IL-17F (or the IgG isotype control) to physiologically aging mice (73-week-old) for 12 weeks (Fig. 5a). We then isolated dermal cells by FACS and performed 10× scRNA-seq, following the previous workflow (see Fig 1a). A total of 16,975 CD45+ cells and 33,262 CD45–/EpCAM– cells were analyzed with the same bioinformatics pipeline as the aged sample analysis.

**Figure 5.**
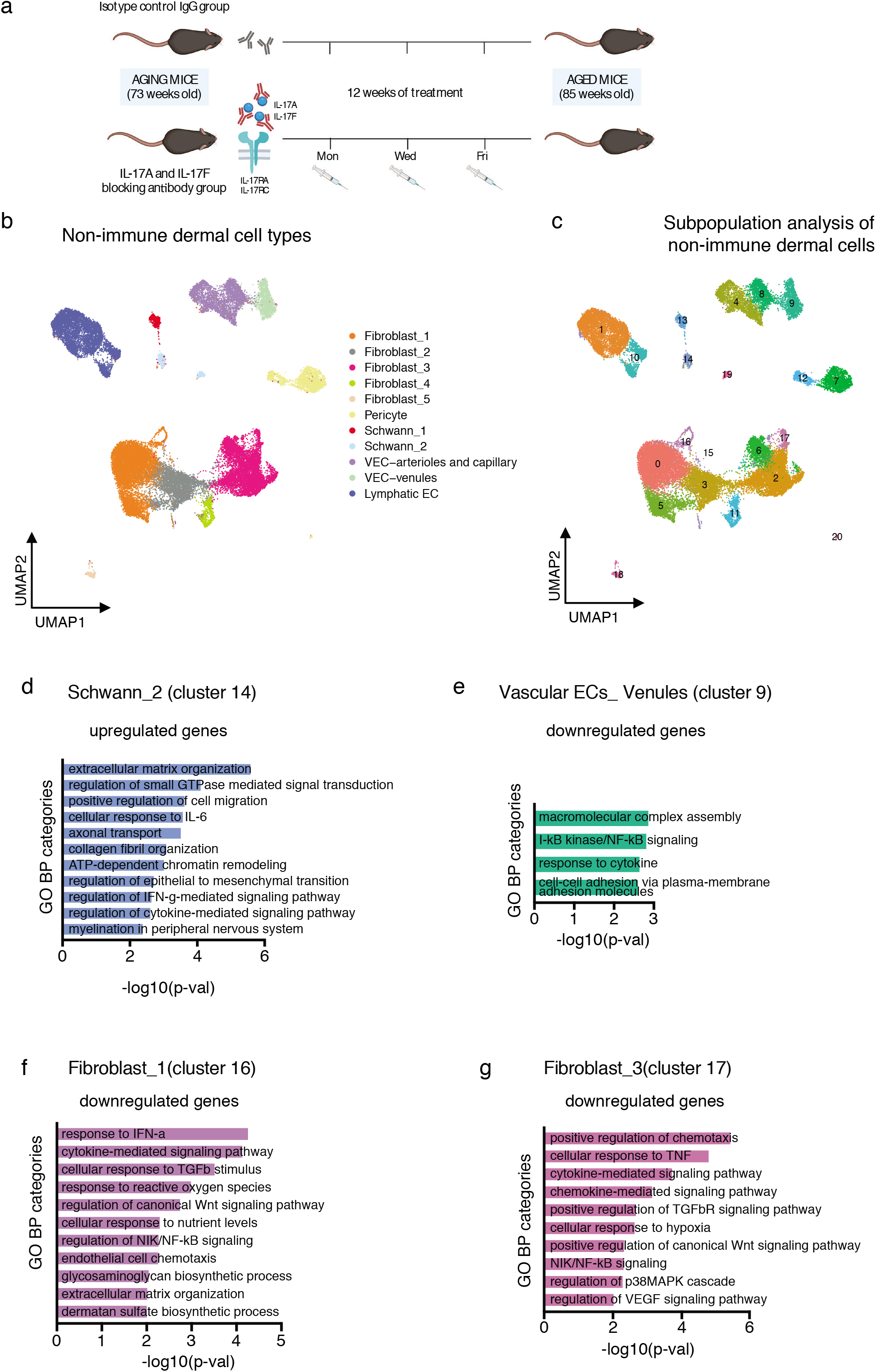
Blockade of IL-17A/F function in aging mice leads to a delay in age-associated traits in non-immune dermal cells. **a,** Diagram showing the anti-IL-17A/F treatment workflow. **b**, UMAP representation of the clusters of non-immune cell types found in the IgG control treated and the anti-IL-17A/F treated dermal preparations. **c**, UMAP representation of the subclusters found in the dermal non-immune population of IgG control treated and anti-IL-17A/F treated mice. For each subgroup of CD45+ cells and CD45–/EpCAM– cells, *n* = 8 mice for the non-aged control group, and *n* = 8 mice for the aged group, with four technical replicates. **d**, Plot of selected GO categories belonging to BP analysis for genes upregulated during IL-17A/F blocking treatment in cluster_14 (belonging to previously described Schwann_2 cells). **e**, Plot of selected GO categories belonging to BP analysis for genes downregulated during IL-17A/F blocking treatment in cluster_9 (belonging to previously described vascular EC-venule cells). **f, g**, Plot of selected GO categories belonging to BP analysis for genes downregulated during IL-17A/F blocking treatment in cluster_16 (**f**) and cluster_17 (**g**) (belonging to the fibroblast_1 and fibroblast_3 clusters, respectively). For **d, e, f** and **g**, the x axis represents –(log_10_) of the *P*-value for each depicted GO category.

#### Non-immune dermal cells

Neutralizing IL-17 did not change the proportions of any non-immune dermal cell type (Supplementary Fig. 3a,b and Supplementary Table 4), and some clusters did not show any significant changes in gene expression upon IL-17 inhibition. However, when we further zoomed into each cell type and carried out a subpopulation analysis, we observed a strong trend towards attenuating age-associated traits related to gene expression in specific cell states (Fig. 5b,c). For example, our previously defined fibroblast_1 cluster could be subclassified into three subclusters, of which only two (cluster_5 and cluster_16) significantly responded to inhibition of IL-17 (Supplementary Table 5).

Our previous single-cell analysis identified two clusters of dermal Schwann cells that corresponded to a more myelinating differentiated state and a progenitor population (Fig. 2a). Interestingly, in the latter, now cluster_14, neutralization of IL-17 led to an upregulation in the expression of genes important for EMT, ECM remodeling and myelination, whose expression was lower in the aged skin (Fig. 5d and Supplementary Tables 5 and 6). Conversely, genes downregulated in Schwann cells during aging, such as *Itga4, Nf1* and some collagen isoforms, were upregulated following IL-17 inhibition.

Of note, blocking IL-17 clearly reverted the trend of aged venule vascular ECs which showed an increased expression of genes related to inflammation (see Fig. 2c), as shown in cluster_9 (Fig. 5e and Supplementary Tables 5 and 6). Strikingly, it also downregulated the expression of genes such as *Cd74*, a key component of the MHCII complex, that we observed to be upregulated in aged ECs (see Fig. 2e). Moreover, *Sele* and *Pecam1*, which constitute markers of vessel activation that favor lymphocyte rolling and extravasation^48^, also were expressed at lower levels following IL-17 inhibition, indicative of a less pro-inflammatory state of the IL-17-neutralized dermal post-capillary venules (Supplementary Tables 5 and 6)..

Aged fibroblasts defined in clusters fibroblast_1 and fibroblast_3 showed an increase in the expression of genes related to inflammation and immune cell recruitment (Fig. 2f,g). Importantly, *in vivo* inhibition of IL-17 prevented these changes as compared to the control IgG–treated aged mouse cohorts, as shown by GO analysis of downregulated genes (Fig. 5f,g, and Supplementary Tables 5 and 6), confirming that fibroblasts are also sensitive to excessive age-related IL-17 signaling.

#### Immune cells

Myeloid cells are responders to IL-17 signaling in inflammatory situations, leading to the activation of the expression of more pro-inflammatory cytokines^64^. Importantly, the upregulation of pro-inflammatory cytokines (such as *Il1b* in clusters macrophage_2, monocyte_1 and dendritic cells) that we observed to be associated to aging was significantly attenuated upon neutralization of IL-17A/F signaling (Supplementary Fig. 3c and Supplementary Tables 5 and 6). We also observed a general attenuation of pro-inflammatory genes in the monocyte_1 cluster (Supplementary Fig. 3d), which we had previously identified as pro-inflammatory monocytes, and whose function was exacerbated during aging.

Altogether, our results indicate that attenuation of IL-17A/F signaling through treatment with neutralizing antibodies pleiotropically attenuated age-related phenotypes in most dermal cell types, with predominant changes observed in pro-inflammatory fibroblasts, vascular ECs and Schwann cells.

### Delay in age-associated traits of epidermal keratinocytes upon IL-17-blockade

We next investigated whether anti-IL-17 treatment could also ameliorate age-associated traits in epidermal keratinocytes. Sustained inflammation in aged epidermis is thought to affect epidermal stem cell fitness, potentially by reducing their regenerative function^2,10,13,17^. In addition, the perturbed communication between epidermal stem cells and dendritic epidermal T cells contributes to delayed wound healing in aged skin^28^.

Bulk RNA-seq of aged and adult epidermal cells confirmed that expression levels of pro-inflammatory cytokines and chemokine signaling increased in aged skin (Supplementary Fig. 4a and Supplementary Table 7). We also observed an upregulation of genes important for other known epidermal aging traits, such as toxic reactive oxygen species (ROS)-related stress ^2^. Importantly, IL-17 blockade led to reduced expression of cytokine-related signaling and chemotactic genes, such as *Ccl5* and *Xcl1*, as well as to downregulation of genes involved in pro-inflammatory and oxidative stress functions (Fig. 6a and Supplementary Table 7). Concomitant to this, there was a reduction in the exacerbated expression of differentiation markers of aged skin, exemplified by lower *Flg* and *Lor* expression, and an increase in the expression of genes related to healthier wound healing, such as *Wnt7a*^65^ (Fig. 6a and Supplementary Table 7).

**Figure 6.**
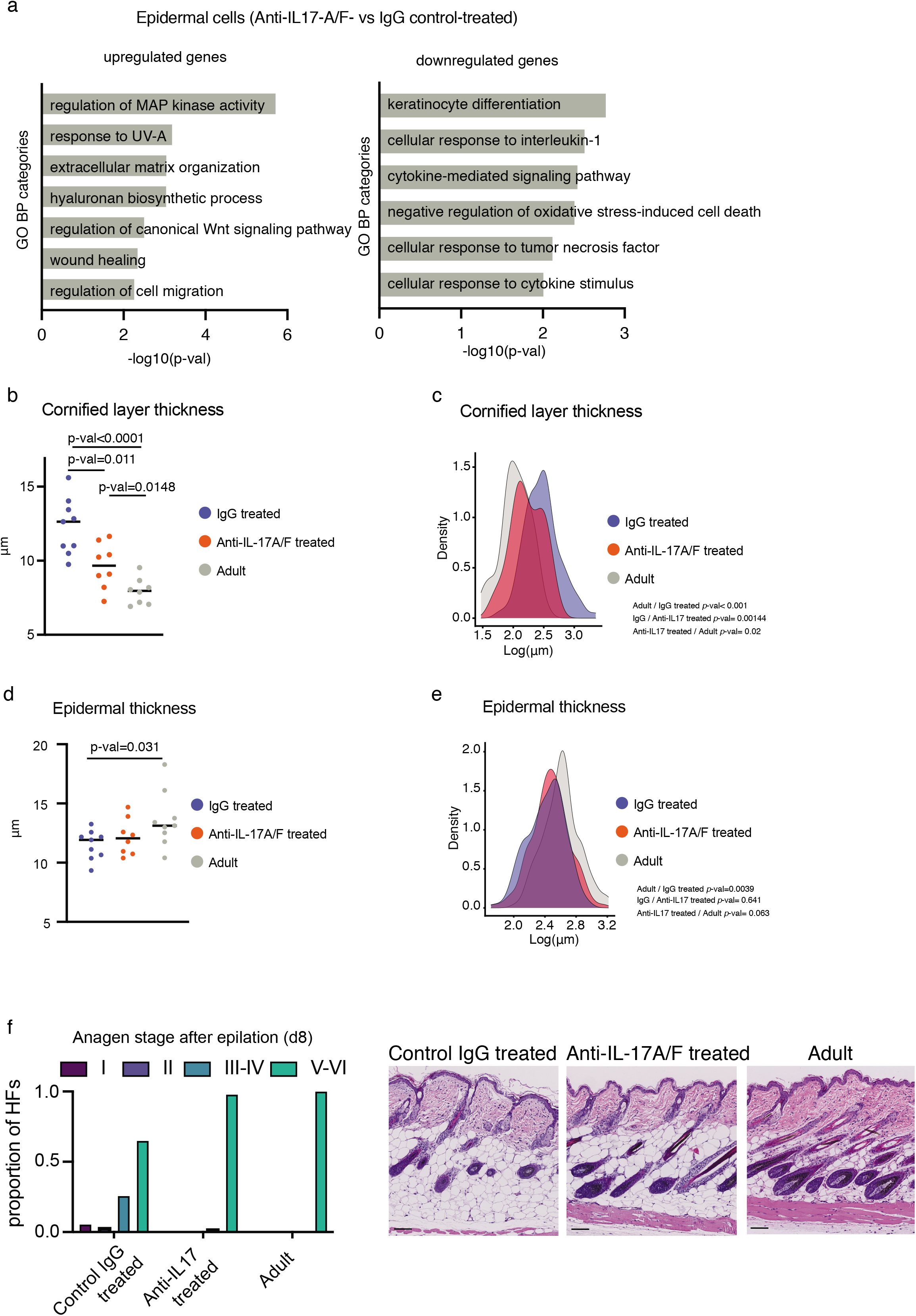
Decreased epidermal age-associated traits upon IL-17A/F blockade. **a**, Plot of selected GO categories belonging to BP analysis for genes upregulated (left panel) and downregulated (right panel) during IL-17A/F blocking treatment in epidermal cells. The x axis represents –(log_10_) of the *P*-value for each depicted GO category. **b**, Dot plot with quantification of the cornified layer thickness (in μm) in adult control (gray), aged treated with control IgG (blue) and aged treated with anti-IL-17A/F (orange). **c**, Histogram showing the distribution of all the values obtained for cornified layer thickness to analyze their distribution; *n* = 8 mice. *P*-values were obtained using mixed linear models and multiple mean comparison using Tukey contrasts. **d**, Dot plot with measures of epidermal thickness (in μm). **e**, Histogram showing the distribution of all the values obtained for epidermal thickness in the 8 mice to analyze their distribution. *P*-values were obtained using mixed linear models and multiple mean comparison by Tukey contrasts. For **b** and **d**, each dot represents the mean of 10 measures (cornified layer) or 20 measures (epidermal thickness) per mouse (*n* = 8 mice) obtained in Hematoxylin and Eosin (H&E) staining of skin sections (see Supplementary Fig. 4b). The line depicts the median of these individual values per age and condition. *P*-values were calculated by Mann Whitney *U* test. **f**, Left, quantification of anagen stages in mouse back skin at 8 days after epilation of i) adult, ii) aged/control IgG, and iii) aged/anti-IL-17A/F antibodies. Right, details of H&E staining of epilated back skins at 8 days post-epilation; bar = 100 μm.

### Youthful skin traits increase after IL-17 signaling is neutralized

A well-defined age-related phenotype in the epidermis is the excessive differentiation program expressed by keratinocytes that leads to an increased thickness of the cornified layer, the outermost impermeable layer that is composed of highly-keratinized, dead enucleated cells^2^. Indeed, the thickness of the cornified layer doubled in the aged mice as compared to the control mice (Fig. 6b,c and Supplementary Fig. 4b). Strikingly, however, *in vivo* systemic inhibition of IL-17 (using anti-IL-17A/F antibodies) on aged mice resulted in a significantly thinner cornified layer, as compared to control (IgG-treated) aged mice (Fig. 6b,c).

We also observed a reduced thickness of the epidermis (which is another hallmark of aging in the skin^9,28,66^) in our mouse cohort (Fig. 6d,e). However, when aged mice were treated with neutralizing antibodies against IL-17A and IL-17F, epidermal thickness significantly increased, as compared to control-treated aged mice (Fig. 6d,e and Supplementary Fig. 4b). Thus, these data suggested that blocking IL-17 signaling could delay the acquisition of age-related traits and decrease the level of inflammation in the epidermis, even when treatment was done in the late adulthood stage of life.

We next asked whether the neutralization of IL-17A/F affected stem cell behavior. Aged hair follicles show a significant lower capacity to enter the growing phase of the hair cycle (anagen)^17^. Anagen can be subdivided into several phases that indicate how far the growth of the hair follicles has advanced^67^. As expected, aged hair follicles in our cohort of aged mice showed a delay in the anagen stage as compared to the adult counterparts at 8 days post-epilation (Fig. 6f). Strikingly, the HFs of the aged/anti-IL-17A/F–treated mice (but not the aged/IgG-treated mice) showed a faster pace in their anagen entry and progression, which was almost identical to adult mice, pointing towards a strong amelioration of the capacity of hair follicle stem cells to become activated as compared to aged IgG-treated control mice (Fig. 6f and Supplementary Fig. 4c). Dermal papilla fibroblasts (which are essential for hair follicle stem cell activation during anagen entry) that are represented in the cluster fibroblast_1 (Fig. 2a), now could be divided into 3 subclusters (cluster_0, cluster_5 and cluster_16) (Fig. 5c). Analyzing the treatment-dependent DEGs in these clusters, we found an upregulation of *Rspo3*, a Wnt agonist whose expression in dermal papilla cells triggers the active phases of hair cycling^68,69^ (Supplementary Fig. 4d). Altogether, we found that aberrant IL-17 signaling was at the hinge between dermal and epidermal cells, and neutralization of it in both cell compartments caused a synergy that worked towards delaying and even ameliorating age-associated skin traits.

## Discussion

During aging, tissue-specific alterations in the niche synergize with stem cell–intrinsic changes to contribute to the development of age-associated traits^17,70-72^. Aging has been proposed to drive a pro-inflammatory microenvironment that perturbs adult stem cell behavior in several tissues, including that of neural, skeletal and hematopoietic stem cells^73–76^. Interestingly, the nature of these signals is highly tissue-dependent and include infiltration of immune cells into the stem cell niche^73^, or a transcriptional switch of the stem cells that create a pro-inflammatory environment that negatively feeds back to their own fitness^74^. Here, we have characterized for the first time, and in an unbiased manner, the effects of the pro-inflammatory cytokine IL-17 on an aging tissue—namely, skin. Our results show that elevated IL-17 signaling, which was specifically secreted by aged dermal CD4+ Th, γδ T and ILCs, orchestrated many of the age-associated tissue dysfunction by exerting pleiotropic effects on all skin compartments (immune and non-immune). IL-17-mediated signaling is heavily linked to the development of chronic inflammatory and autoimmune diseases^54,58,77,78^. In skin, these diseases include psoriasis, pemphigous and alopecia areata^55^. Even if none of the clinical signs of these diseases are common with physiological aging, they share an increased aberrant IL-17-based signaling that impedes correct skin function. Alopecia areata is an autoimmune disease characterized by loss of hair in defined areas^79,80^. This could point towards a relationship between IL-17 and HF cycling. Indeed, conflicting reports of psoriatic patients undergoing IL-17-neutralizing treatments show that they frequently develop either hypertrichosis or alopecia (e.g., excessive hair growth or loss, respectively), pointing out that blocking an excessive IL-17 signaling could influence hair follicle cycling in humans^81,82^. On the other hand, the excessive IL-17 signaling observed in aged dermal cells might underlie the development of bullous pemphigoid, an autoimmune disorder with aberrant IL-17 signaling that is characterized by skin blistering, and whose incidence is increased in elderly people^53^. Our results strongly suggest that the local environment of the aged skin intriguingly resembles a low-level but persistent state of chronic inflammation that is reminiscent of that in serious skin diseases. We hypothesize that treatments with biologicals already approved against psoriasis or pemphigus could also be useful for other age-associated ailments, such as the inability to repair damaged skin that is exemplified by chronic wounds in elderly.

## Acknowledgments

Research in the S.A.B. lab is supported partially by the European Research Council (ERC) under the European Union’s Horizon 2020 research and innovation programme (Grant agreement No. 787041), the Government of Cataluña (SGR grant), the Government of Spain (MINECO), the La Marató/TV3 Foundation, the Foundation Lilliane Bettencourt, the Spanish Association for Cancer Research (AECC) and The Worldwide Cancer Research Foundation (WCRF). This work was partially supported by a Leonardo grant (BBVA Foundation) awarded to G.S. P.S. was awarded by a Spanish Ministry of Economy and Development fellowship (ID BES-2017-081279). M.E and E.M. were supported by Ministerio de Ciencia e Innovación (MCI), Agencia Estatal de Investigación (AEI) and the European Development Regional Fund, ‘A way to make Europe’ ERDF (RTI2018-094049-B-I00). H.H. has received funding from the Ministerio de Ciencia, Innovación y Universidades (SAF2017-89109-P; AEI/FEDER, UE). The IRB Barcelona is a Severo Ochoa Center of Excellence (MINECO award SEV-2015-0505). We would like to thank the histology and genomics facilities of the IRB Barcelona for their assistance in this work. We would like to thank the genomics unit at the CRG for assistance with bulk RNA sequencing. We thank Veronica Raker for manuscript editing.

## Author contributions

P.S., J.B. and G.S. designed the experiments, collected, analyzed and interpreted the data. E.M. and M.C. analyzed and interpreted the scRNA-seq data. O.R. and E.B. performed the RNA-seq bioinformatics analysis. M.E., H.H. and L.D. supported financially the project and helped with data interpretation. G.S. and S.A.B. conceived and supervised the project and drafted the article. All authors contributed to the final manuscript.

## Methods

### Mouse handling and husbandry

Mice were housed under a regimen of 12-h/12-h light/dark cycles and specific pathogen free conditions. All procedures were evaluated and approved by the Ethical Committee for Animal Experimentation (CEEA) from the Government of Catalunya. Retired C57Bl/6J breeder females were purchased from Charles River and were aged until the desired age in the animal facility at the Barcelona Science Park (PCB). Control adult mice were either bred in-house or purchased to Charles River to generate matching cohorts. Mostly female mice were used due to this fact, and to differences between skin digesting efficiency between sexes, as longer male skin digestion times are required and this reduced the survival of sorted dermal cells, skewing the results towards the most resilient cell types. Mice were always sacrificed in the dark period, to coincide with their active phases. Old mice were between 80- to 90-weeks of age, and adult mice were between 17- to 25-weeks of age.

### In vivo anti-IL-17A/F neutralizing treatment

A cohort of aging (73-week-old) mice was randomly distributed into two groups, and these were treated with either i) a mixture of 105 μg anti-IL-17A (clone 17F3, BioXCell) and 105 μg anti-IL-17F (clone MM17F8F5.1A9, BioXCell), or ii) 210 μg control IgG1 (clone MOPC-2, BioXCell). Injections of 100 μl antibodies were administered intraperitoneally (i.p.) and performed 3 times per week at the same time of the day (at 2- to 3-h into the dark phase). After 12 weeks of treatment, mice were either sacrificed to obtain samples for dermal cells for 10× scRNA-seq, bulk RNA-seq of the epidermis and histology, or they were used for epilation.

### Epilation

Mice were anesthetized using a mixture of ketamine (75 mg/kg body weight) and medetomidine (1 mg/kg) via i.p. injection. Buprenorphine (0.05 mg/kg) was injected subcutaneously as a pain killer and an anti-inflammatory treatment. An area of about 2- to 3-cm^2^ of back skin was epilated using wax papers until no hair was visible (usually 2 or 3 rounds were enough). Atipamezole (1 mg/kg) was injected to revert anesthesia, and mice were left on a warm blanket to recover. Afterwards, mice were housed individually to avoid contact of the epilated areas. Animal health status was monitored daily. At 8 days after hair removal, mice were sacrificed, and images of shaved back skin were taken. Samples of small portions of back skin were fixed in neutral buffered formalin (10%) for 3 h at room temperature, dehydrated, and included in paraffin blocks for later histological assessment.

### Epidermal and dermal cell isolation

Mice were sacrificed, and whole torso skin was removed as fast as possible. Hypodermal fat was removed with scalpel and after 2 washes in PBS, skins were floated (dermis-side down) in a Dispase II solution (5 mg/mL; D4693 Sigma-Aldrich) in PBS for 30- to 40 min at 37°C. Epidermises were removed with a scalpel. For dermal cell isolation: dermises were mechanically dissociated using a McIlwain Tissue Chopper (The Mickle Laboratory Engineering Co. LTD) and then further digested in Liberase TM (6.5 Wünsch units/reaction, Roche) diluted in DMEM (41965, Thermo Fisher Scientific) for 20 to 30 min at 37°C with gentle agitation. Afterwards, DNase I (1 mg/ml DN25; Sigma-Aldrich) was added to the mix and incubated for 15 min at 37°C without agitation. Digested dermises were strained first through a 100-μm strainer and then through a 40-μm strainer, to obtain single-cell suspensions. For epidermal cell isolation: epidermises were removed with a scalpel and mechanically dissociated using a McIlwain Tissue Chopper (The Mickle Laboratory Engineering Co. LTD). They were then strained through a 100-μm and then a 40-μm strainer to obtain single-cell suspensions. and frozen in 1 ml of TRIzol (Invitrogen) for posterior RNA isolation. RNA was extracted from epidermal cell pellets frozen in TRIzol using the RNeasy Mini Kit (Qiagen) and further processed for mRNA-seq with the Illumina sequencing technology.

### Library construction and sequencing

For the adult *vs* aged comparison, libraries were prepared using the TruSeq Stranded Total RNA Library Prep Kit with Ribo-Zero Human/Mouse/Rat Kit (cat. RS-122-2201/2202, Illumina) according to the manufacturer’s protocol, using 150 to 300 ng of total RNA; ribosomal RNA depletion, RNA was then fragmented for 4.5 min at 94°C. The remaining steps were followed according to the manufacturer’s instructions. Final libraries were analyzed on an Agilent Technologies 2100 Bioanalyzer system using the Agilent DNA 1000 chip to estimate the quantity and validate the size distribution; libraries were then quantified by qPCR using the KAPA Library Quantification Kit KK4835 (cat. 07960204001, Roche) prior to amplification with Illumina’s cBot. Finally, libraries were sequenced on the Illumina HiSeq 2500 sequencing system using paired-end 50-base pair (bp)-long reads.

For the IL-17 neutralized *vs* control samples, libraries were prepared using the TruSeq stranded mRNA Library Prep (cat. 20020595, Illumina) according to the manufacturer’s protocol, to convert total RNA into a library of template molecules of known strand origin that is suitable for subsequent cluster generation and DNA sequencing. Briefly, 500–1000 ng total RNA was used for poly(A)-mRNA selection using poly-T oligonucleotides attached to magnetic beads with two rounds of purification. During the second elution of the poly-A RNA, RNA was fragmented under elevated temperature and primed with random hexamers for cDNA synthesis. The cleaved RNA fragments were then copied into first-strand cDNA using reverse transcriptase (SuperScript II, ref. 18064-014, Invitrogen) and random primers. Note that addition of actinomycin D to the First Stand Synthesis Act D mix (FSA) improves strand specificity by preventing spurious DNA-dependent synthesis while allowing RNA-dependent synthesis. Second-strand cDNA was then synthesized by removing the RNA template and synthesizing a replacement strand, incorporating dUTP in place of dTTP to generate ds cDNA using DNA polymerase I and RNase H. These cDNA fragments then had a single A base added to the 3’-ends of the blunt fragments, to prevent them from ligating to one another during the adapter ligation. A corresponding single T-nucleotide on the 3’-end of the adapter provided a complementary overhang for ligating the adapter to the fragments, ensuring a low rate of chimera (concatenated template) formation. Subsequent ligation of the multiple indexing adapter to the ends of the double-strand cDNA was carried out. Finally, PCR selectively enriched DNA fragments with adapter molecules on both ends, and the amount of DNA in the library was amplified. PCR was performed with a PCR Primer Cocktail that anneals to the ends of the adapters. Final libraries were analyzed using Bioanalyzer DNA 1000 or Fragment Analyzer Standard Sensitivity (Agilent) to estimate the quantity and validate the size distribution; libraries were then quantified by qPCR using the KAPA Library Quantification Kit KK4835 (Roche) prior to amplification with Illumina’s cBot. Finally, libraries were sequenced on the Illumina HiSeq 2500 sequencing system using single-end 50-bp-long reads.

### Hematoxylin and eosin staining

Paraffin blocks were cut in 4-μm sections. Hematoxylin and eosin (H&E) staining was performed according to the standard protocol. Images were acquired using a NanoZoomer scanner (Hamamatsu) at 20× magnification. Scaled images were analyzed with Qupath v0.3.0, with the thicknesses measured. All values per mouse were averaged, and graphical representations were performed with GraphPad Prism 9. Eight mice per condition and age were used.

### Flow cytometry and cell sorting

(giving CD45+, CD45–/EpCAM– and CD31+/MHCII) For 10× scRNA-seq, single-cell dermal suspensions were incubated with CD45–APC (clone 30-F11, 1:100, BD Biosciences) and EpCAM–PE (clone G8.8, 1:200, BD Biosciences) for 45 min on ice. After two washes in PBS, cells were resuspended in 2 μg/ml DAPI (32670, Sigma-Aldrich) to stain DNA and analysed using a BD FACSAria Fusion flow cytometer, in which CD45+/− cells were obtained and EpCAM+ cells excluded.

For MHCII analysis in CD31+ cells, single-cell dermal suspensions were incubated with CD31-PE (clone MEC13.3, 1:100, BD Bioscience) MHCII-BV650 (I-A/I-E, clone M5/114.15.2, 1:500, BioLegend), LYVE1-eFluor 660 (clone ALY7, 1:100, Invitrogen), and (as exclusion markers) CD45–FITC (clone 30-F11, 1:100, eBioscience), CD117-FITC (clone 2B8, 1:100, BDBiosciences), CD41-FITC (clone eBioMWReg30, 1:100, Invitrogen), and EpCAM–FITC (clone G8.8, 1:100, Biolegend). After two washes with PBS, cells were resuspended in 2 ug/ml DAPI (32670, Sigma-Aldrich) to stain DNA and analysed using a BD FACSAria Fusion flow cytometer.

### 10× scRNA-seq

Stained dermal single-cell suspensions were prepared as explained above. CD45+ and CD45–/EpCAM– cells were FACS sorted in a BD Fusion cell sorter separately to enrich for the less abundant CD45+ cells, following the sorting strategy depicted in Suppl. Fig. 1a. Cells were collected in PBS + 0.5% BSA at 4°C in LoBind tubes (Eppendorf) and processed immediately with the microfluidics Chromium platform (10**×** Genomics).

### Data pre-processing

Sequences were demultiplexed and aligned according to Cell Ranger pipeline (version 6.0.0) with default parameters. Sequencing reads were mapped against the mouse GRCm38 reference genome to generate feature-barcode matrixes, separately for all CD45+ and CD45 –replicates.

### QC and technical bias corrections

Gene count matrixes were analyzed with Seurat package (version 4.0.4) in R (version 4.0.3)^83^. The replicates were merged, analyzed, and annotated separately for CD45+ and CD45– datasets before their integration. Cells were filtered with more than 10% of mitochondrial gene content and genes not found in at least 5 cells. As part of the quality control, cells situated between the minimum and 1st quartile (according to the distribution of number of genes per cell of each compartment dataset) were removed. To avoid contamination of epithelial cells in the immune compartment, EPCAM+ cells found in CD45+ sorted cells were filtered out. Additionally, to remove the technical biases from merging replicates within CD45+, the differentially expressed (DE) genes between the replicates were computed using the *FindAllMarkers* function, and overlapping genes between the top 500 DE genes and the highly variable genes (HVGs) were removed before clustering.

### Clustering

Cell-to-cell variations were normalized by the expression values using a scale factor of 100,000 and a log transformation. The gene expression measurements were scaled and centered. The scaled Z-score values were then used as normalized gene measurement input for clustering and for visualizing differences in expression between cell clusters. HVGs were selected by assessing the relationship of log(variance) and log(mean) and choosing those with highest variance-to-mean ratio. Principal components analysis (PCA) was used to reduce the dimensionality of the dataset, and *ElbowGraph* was used to select the number of dimensions for the clustering for significant principal components. Cluster identification was performed using the functions *FindNeighbors* and *FindClusters*, which calculates the k-nearest neighbors and generates the shared nearest neighbor (SNN) graph to cluster the cells. The algorithm applied was the Louvain method, which allows the number of clusters to be tuned with a resolution parameter. To explore the clusters in more detail, the resolution parameter was increased in *FindClusters* function or *FindSubCluster* was used for specific clusters. The Uniform Manifold Approximation and Projection (UMAP) was used as a non-dimensional reduction method, to visualize the clustering.

### Cell type annotation

To annotate the cell types and states, enriched genes were first identified in each of the clusters using *FindAllMarkers* function, using the Wilcoxon Rank Sum test to find cluster-specific markers. These cluster-specific genes were then explored to find previously reported cell population marker genes. Some examples of marker genes used to annotate the cell populations are given here: Cd4+ T cells: *Cd28, Cd4*; Cd8+ T cells: *Cd8a, Cd8b1*; dermal cells (DC): *Cd207;* fibroblast_1: *Crabp1, Inhba, Notum*; fibroblast_2: *Col1a1, Col1a2, Cd34, Robo1, Col3a1*; fibroblast_3: *Efemp1, Il1r2, Ccl11*; fibroblast_4: *Col11a1, Aspn, Coch*; fibroblast_5: *Myoc, Dcn; ILC: Il13, Kit*; lymphatic endothelial cells (ECs): *Lyve1, Hes1*; macrophage_1: *Il1b;* macrophage_2: *Tnfsf9*; macrophage_3: *Ear2, Cd163;* monocyte_1: *Plac8, Cd14*; monocyte_2: *Ccl8, C1qa;* monocyte_3*: Retnla, C1qb, Ccr2*; NK cells: *Gzmc, Ccl5, Nkg7*; pericytes: *Acta2, Rgs5*; proliferating macrophages: *Mki67*; proliferating T cells: *Hmgb2proliferating T cell*; Schwann cells: *Cryab, Plekha4, Scn7a*; Schwann cells_1: *Kcna1*; Schwann cells_2: *Sox10*; T regulatory cells*: Foxp3;* VEC arterioles and capillary: *Ptprb, Flt1*; VEC venules: *Aqp1, Sele, Pecam1*; and γδ T cells: *Trdc, Tcrg-C1*.

### Differential expression (DE) analysis for each cluster

To find differentially expressed genes between adult *vs*. aged, and IgG treated *vs*. anti-IL-17A/F treated, in the different annotated cell type populations, DE analysis was performed between conditions for each cluster with *FindMarkers* function.

### Age effect analysis

The effects of aging on dermis were evaluated based on how cell types differ transcriptionally between adult and aged mice. For this, the similarities of cell type– specific markers were evaluated by comparing the Jaccard Indexes between age groups across cell types. The Jaccard Index was computed using MatchScore2, which considers the top 100 DE markers as a signature of each population^84^. To further assess the age-related effects among the immune cells, a deep-learning approach was created that is based on auto-encoders. During training, this method can identify features that are relevant for the data structure and then use them to predict the different cell types in a test dataset. To generate the model, adult data was divided into two smaller subsets that could be used as training and test sets. The model displayed high probability scores (*P* > 0.5) in the prediction of cell types from the test set among the different immune cell types and a low rate of unpredicted cells (unclassified rate < 5%). If the rate of unclassified cell type is strictly biased for aged cells, this would reflect the effect of aging on the cell phenotype. Based on this assumption, an ***age deviance score*** was defined as 1-q, whereby q is the probability of the unpredicted cell being in the true corresponding cell type class in aged cells. The proportion of unclassified cells within each cell type in aged cells, and the proportion of age-affected cells, were then measured and was normalized by the model’s error rate (e.g., by subtracting the cell proportion in each adult cell type that could not be predicted by the model).

### Bioinformatics analyses of bulk RNA-seq data

FastQ files were aligned against the mm10 reference genome using STAR 2.5.2b^85^ using default options. Unless otherwise specified, all downstream analyses were performed using R 3.5.1. Differentially expressed genes (DEGs) between conditions were determined using DESeq2 1.22.1^86^, using mm10 gene counts as generated with the featureCounts function from the RSubread package version 1.32.4^87^ with options: *annot.inbuilt=‘mm10’,allowMultiOverlap=TRUE,countMultiMappingReads=FALSE,minMQS=1,ignoreDup=FALSE*). Genes were selected as DEGs using the thresholds |1fcShrink foldChange|>1.25 and Benjamini-Hochberg–determined *P* < 0.1, using experimental batch as covariate. Gene set enrichment analysis was performed using gene set collections at *Mus musculus* gene symbol level. The gene set collections used were: GOBP, GOMF, GOCC, and KEGG, obtained using the org.Mm.eg.db package, November 2014; GOSLIM, obtained from geneontology.org, November 2014; and Broad Hallmarks, obtained from the Broad Institute MSigDB website (https://www.gsea-msigdb.org/gsea/msigdb/) and mapped from human to mouse genes using homology information from Ensembl biomart archive July 2016. Analyses were performed using regularized log transformation (rlog) applied to the count data using the DESeq2 R package 1.22, with ROAST^88^ and the <u>MaxMean</u> statistic (http://statweb.stanford.edu/~tibs/GSA/).

### GO analysis

GO analysis was performed with enrichR^89^ (https://maayanlab.cloud/Enrichr/). A category was considered significant if *P* < 0.01.

### Statistical analysis

Generally, graphs show median and *P*-values were obtained with the Mann-Whitney *U* test with Prism v9, unless otherwise stated in the figure legend. Statistical analyses of epidermal skin and cornified layer thickness were performed using mixed linear models in R3.5.1 with lme4 library 1.1-23 and multcomp 1.4-9.

### Data availability

10X Single cell sequencing data is deposited at the NCBI GEO repository, accession GSE193920.

Bulk RNA-seq data is deposited at the NCBI GEO repository, accessions GSE190182 and GSE190393.

## References

1 Sato, S. et al. Circadian Reprogramming in the Liver Identifies Metabolic Pathways of Aging. Cell 170, 664–677 e611, doi:10.1016/j.cell.2017.07.042 (2017).

2 Solanas, G. et al. Aged Stem Cells Reprogram Their Daily Rhythmic Functions to Adapt to Stress. Cell 170, 678–692 e620, doi:10.1016/j.cell.2017.07.035 (2017).

3 Lopez-Otin, C., Blasco, M. A., Partridge, L., Serrano, M. & Kroemer, G. The hallmarks of aging. Cell 153, 1194–1217, doi:10.1016/j.cell.2013.05.039 (2013).

4 Green, D. R., Galluzzi, L. & Kroemer, G. Mitochondria and the autophagyinflammation-cell death axis in organismal aging. Science 333, 1109–1112, doi:10.1126/science.1201940 (2011).

5 Dimri, G. P. et al. A biomarker that identifies senescent human cells in culture and in aging skin in vivo. Proc Natl Acad Sci U S A 92, 9363–9367, doi:10.1073/pnas.92.20.9363 (1995).

6 Salvioli, S. et al. Immune system, cell senescence, aging and longevity--inflamm-aging reappraised. Curr Pharm Des 19, 1675–1679 (2013).

7 Solanas, G. & Benitah, S. A. Regenerating the skin: a task for the heterogeneous stem cell pool and surrounding niche. Nat Rev Mol Cell Biol 14, 737–748, doi:10.1038/nrm3675 (2013).

8 Ghadially, R., Brown, B. E., Sequeira-Martin, S. M., Feingold, K. R. & Elias, P. M. The aged epidermal permeability barrier. Structural, functional, and lipid biochemical abnormalities in humans and a senescent murine model. J Clin Invest 95, 2281–2290, doi:10.1172/JCI117919 (1995).

9 Giangreco, A., Qin, M., Pintar, J. E. & Watt, F. M. Epidermal stem cells are retained in vivo throughout skin aging. Aging Cell 7, 250–259, doi:10.1111/j.1474-9726.2008.00372.x (2008).

10 Doles, J., Storer, M., Cozzuto, L., Roma, G. & Keyes, W. M. Age-associated inflammation inhibits epidermal stem cell function. Genes Dev 26, 2144–2153, doi:10.1101/gad.192294.112 (2012).

11 Benitah, S. A. & Welz, P. S. Circadian Regulation of Adult Stem Cell Homeostasis and Aging. Cell Stem Cell 26, 817–831, doi:10.1016/j.stem.2020.05.002 (2020).

12 Choi, E. H. Aging of the skin barrier. Clin Dermatol 37, 336–345, doi:10.1016/j.clindermatol.2019.04.009 (2019).

13 Ge, Y. et al. The aging skin microenvironment dictates stem cell behavior. Proc Natl Acad Sci U S A 117, 5339–5350, doi:10.1073/pnas.1901720117 (2020).

14 Liu, N. et al. Stem cell competition orchestrates skin homeostasis and ageing. Nature 568, 344–350, doi:10.1038/s41586-019-1085-7 (2019).

15 Matsumura, H. et al. Hair follicle aging is driven by transepidermal elimination of stem cells via COL17A1 proteolysis. Science 351, aad4395, doi:10.1126/science.aad4395 (2016).

16 Giangreco, A., Goldie, S. J., Failla, V., Saintigny, G. & Watt, F. M. Human skin aging is associated with reduced expression of the stem cell markers beta1 integrin and MCSP. J Invest Dermatol 130, 604–608, doi:10.1038/jid.2009.297 (2010).

17 Koester, J. et al. Niche stiffening compromises hair follicle stem cell potential during ageing by reducing bivalent promoter accessibility. Nat Cell Biol 23, 771–781, doi:10.1038/s41556-021-00705-x (2021).

18 Mahmoudi, S. et al. Heterogeneity in old fibroblasts is linked to variability in reprogramming and wound healing. Nature 574, 553–558, doi:10.1038/s41586-019-1658-5 (2019).

19 Gould, L. et al. Chronic wound repair and healing in older adults: current status and future research. J Am Geriatr Soc 63, 427–438, doi:10.1111/jgs.13332 (2015).

20 Norman, R. A. Geriatric dermatology. Dermatol Ther 16, 260–268, doi:10.1046/j.1529-8019.2003.01636.x (2003).

21 Jin, S. P. et al. Changes in tight junction protein expression in intrinsic aging and photoaging in human skin in vivo. J Dermatol Sci 84, 99–101, doi:10.1016/j.jdermsci.2016.07.002 (2016).

22 Salzer, M. C. et al. Identity Noise and Adipogenic Traits Characterize Dermal Fibroblast Aging. Cell 175, 1575–1590 e1522, doi:10.1016/j.cell.2018.10.012 (2018).

23 Sole-Boldo, L. et al. Single-cell transcriptomes of the human skin reveal age-related loss of fibroblast priming. Commun Biol 3, 188, doi:10.1038/s42003-020-0922-4 (2020).

24 Joost, S. et al. The Molecular Anatomy of Mouse Skin during Hair Growth and Rest. Cell Stem Cell 26, 441–457 e447, doi:10.1016/j.stem.2020.01.012 (2020).

25 Reynolds, G. et al. Developmental cell programs are co-opted in inflammatory skin disease. Science 371, doi:10.1126/science.aba6500 (2021).

26 Tabula Muris, C. A single-cell transcriptomic atlas characterizes ageing tissues in the mouse. Nature 583, 590–595, doi:10.1038/s41586-020-2496-1 (2020).

27 Dubois, A., Gopee, N., Olabi, B. & Haniffa, M. Defining the Skin Cellular Community Using Single-Cell Genomics to Advance Precision Medicine. J Invest Dermatol 141, 255–264, doi:10.1016/j.jid.2020.05.104 (2021).

28 Keyes, B. E. et al. Impaired Epidermal to Dendritic T Cell Signaling Slows Wound Repair in Aged Skin. Cell 167, 1323–1338 e1314, doi:10.1016/j.cell.2016.10.052 (2016).

29 Ali, N. et al. Regulatory T Cells in Skin Facilitate Epithelial Stem Cell Differentiation. Cell 169, 1119–1129 e1111, doi:10.1016/j.cell.2017.05.002 (2017).

30 Bhushan, M. et al. Tumour necrosis factor-alpha-induced migration of human Langerhans cells: the influence of ageing. Br J Dermatol 146, 32–40, doi:10.1046/j.1365-2133.2002.04549.x (2002).

31 Pilkington, S. M. et al. Lower levels of interleukin-1beta gene expression are associated with impaired Langerhans’ cell migration in aged human skin. Immunology 153, 60–70, doi:10.1111/imm.12810 (2018).

32 Chen, X. et al. IL-17R-EGFR axis links wound healing to tumorigenesis in Lrig1(+) stem cells. J Exp Med 216, 195–214, doi:10.1084/jem.20171849 (2019).

33 Stuart, T. et al. Comprehensive Integration of Single-Cell Data. Cell 177, 1888–1902 e1821, doi:10.1016/j.cell.2019.05.031 (2019).

34 Zou, Z. et al. A Single-Cell Transcriptomic Atlas of Human Skin Aging. Dev Cell 56, 383–397 e388, doi:10.1016/j.devcel.2020.11.002 (2021).

35 Muhl, L. et al. Single-cell analysis uncovers fibroblast heterogeneity and criteria for fibroblast and mural cell identification and discrimination. Nat Commun 11, 3953, doi:10.1038/s41467-020-17740-1 (2020).

36 Rendl, M., Polak, L. & Fuchs, E. BMP signaling in dermal papilla cells is required for their hair follicle-inductive properties. Genes Dev 22, 543–557, doi:10.1101/gad.1614408 (2008).

37 Jessen, K. R. & Arthur-Farraj, P. Repair Schwann cell update: Adaptive reprogramming, EMT, and stemness in regenerating nerves. Glia 67, 421–437, doi:10.1002/glia.23532 (2019).

38 Balakrishnan, A. et al. Insights Into the Role and Potential of Schwann Cells for Peripheral Nerve Repair From Studies of Development and Injury. Front Mol Neurosci 13, 608442, doi:10.3389/fnmol.2020.608442 (2020).

39 Carr, M. J. & Johnston, A. P. Schwann cells as drivers of tissue repair and regeneration. Curr Opin Neurobiol 47, 52–57, doi:10.1016/j.conb.2017.09.003 (2017).

40 Parfejevs, V. et al. Injury-activated glial cells promote wound healing of the adult skin in mice. Nat Commun 9, 236, doi:10.1038/s41467-017-01488-2 (2018).

41 Clements, M. P. et al. The Wound Microenvironment Reprograms Schwann Cells to Invasive Mesenchymal-like Cells to Drive Peripheral Nerve Regeneration. Neuron 96, 98–114 e117, doi:10.1016/j.neuron.2017.09.008 (2017).

42 Arthur-Farraj, P. J. et al. Changes in the Coding and Non-coding Transcriptome and DNA Methylome that Define the Schwann Cell Repair Phenotype after Nerve Injury. Cell Rep 20, 2719–2734, doi:10.1016/j.celrep.2017.08.064 (2017).

43 Abdo, H. et al. Specialized cutaneous Schwann cells initiate pain sensation. Science 365, 695–699, doi:10.1126/science.aax6452 (2019).

44 Buttner, R. et al. Inflammaging impairs peripheral nerve maintenance and regeneration. Aging Cell 17, e12833, doi:10.1111/acel.12833 (2018).

45 Painter, M. W. et al. Diminished Schwann cell repair responses underlie age-associated impaired axonal regeneration. Neuron 83, 331–343, doi:10.1016/j.neuron.2014.06.016 (2014).

46 Bentov, I. & Reed, M. J. The effect of aging on the cutaneous microvasculature. Microvasc Res 100, 25–31, doi:10.1016/j.mvr.2015.04.004 (2015).

47 Mai, J., Virtue, A., Shen, J., Wang, H. & Yang, X. F. An evolving new paradigm: endothelial cells--conditional innate immune cells. J Hematol Oncol 6, 61, doi:10.1186/1756-8722-6-61 (2013).

48 Pober, J. S. & Sessa, W. C. Evolving functions of endothelial cells in inflammation. Nat Rev Immunol 7, 803–815, doi:10.1038/nri2171 (2007).

49 Laina, A., Stellos, K. & Stamatelopoulos, K. Vascular ageing: Underlying mechanisms and clinical implications. Exp Gerontol 109, 16–30, doi:10.1016/j.exger.2017.06.007 (2018).

50 Pasparakis, M., Haase, I. & Nestle, F. O. Mechanisms regulating skin immunity and inflammation. Nat Rev Immunol 14, 289–301, doi:10.1038/nri3646 (2014).

51 Eming, S. A., Martin, P. & Tomic-Canic, M. Wound repair and regeneration: mechanisms, signaling, and translation. Sci Transl Med 6, 265sr266, doi:10.1126/scitranslmed.3009337 (2014).

52 Li, X., Bechara, R., Zhao, J., McGeachy, M. J. & Gaffen, S. L. IL-17 receptor-based signaling and implications for disease. Nat Immunol 20, 1594–1602, doi:10.1038/s41590-019-0514-y (2019).

53 Chakievska, L. et al. IL-17A is functionally relevant and a potential therapeutic target in bullous pemphigoid. J Autoimmun 96, 104–112, doi:10.1016/j.jaut.2018.09.003 (2019).

54 Blauvelt, A. & Chiricozzi, A. The Immunologic Role of IL-17 in Psoriasis and Psoriatic Arthritis Pathogenesis. Clin Rev Allergy Immunol 55, 379–390, doi:10.1007/s12016-018-8702-3 (2018).

55 Liu, T. et al. The IL-23/IL-17 Pathway in Inflammatory Skin Diseases: From Bench to Bedside. Front Immunol 11, 594735, doi:10.3389/fimmu.2020.594735 (2020).

56 Warren, R. B. et al. Bimekizumab versus Adalimumab in Plaque Psoriasis. N Engl J Med 385, 130–141, doi:10.1056/NEJMoa2102388 (2021).

57 Chen, H. C. et al. IL-7-dependent compositional changes within the gammadelta T cell pool in lymph nodes during ageing lead to an unbalanced anti-tumour response. EMBO Rep 20, e47379, doi:10.15252/embr.201847379 (2019).

58 Ouyang, X. et al. Potentiation of Th17 cytokines in aging process contributes to the development of colitis. Cell Immunol 266, 208–217, doi:10.1016/j.cellimm.2010.10.007 (2011).

59 Bharath, L. P. et al. Metformin Enhances Autophagy and Normalizes Mitochondrial Function to Alleviate Aging-Associated Inflammation. Cell Metab 32, 44–55 e46, doi:10.1016/j.cmet.2020.04.015 (2020).

60 Ciofani, M. et al. A validated regulatory network for Th17 cell specification. Cell 151, 289–303, doi:10.1016/j.cell.2012.09.016 (2012).

61 Yamazaki, T. et al. CCR6 regulates the migration of inflammatory and regulatory T cells. J Immunol 181, 8391–8401, doi:10.4049/jimmunol.181.12.8391 (2008).

62 Carr, T. M., Wheaton, J. D., Houtz, G. M. & Ciofani, M. JunB promotes Th17 cell identity and restrains alternative CD4(+) T-cell programs during inflammation. Nat Commun 8, 301, doi:10.1038/s41467-017-00380-3 (2017).

63 Kobayashi, T., Ricardo-Gonzalez, R. R. & Moro, K. Skin-Resident Innate Lymphoid Cells - Cutaneous Innate Guardians and Regulators. Trends Immunol 41, 100–112, doi:10.1016/j.it.2019.12.004 (2020).

64 McGinley, A. M. et al. Interleukin-17A Serves a Priming Role in Autoimmunity by Recruiting IL-1beta-Producing Myeloid Cells that Promote Pathogenic T Cells. Immunity 52, 342–356 e346, doi:10.1016/j.immuni.2020.01.002 (2020).

65 Ito, M. et al. Wnt-dependent de novo hair follicle regeneration in adult mouse skin after wounding. Nature 447, 316–320, doi:10.1038/nature05766 (2007).

66 Branchet, M. C., Boisnic, S., Frances, C. & Robert, A. M. Skin thickness changes in normal aging skin. Gerontology 36, 28–35, doi:10.1159/000213172 (1990).

67 Muller-Rover, S. et al. A comprehensive guide for the accurate classification of murine hair follicles in distinct hair cycle stages. J Invest Dermatol 117, 3–15, doi:10.1046/j.0022-202x.2001.01377.x (2001).

68 Harshuk-Shabso, S., Dressler, H., Niehrs, C., Aamar, E. & Enshell-Seijffers, D. Fgf and Wnt signaling interaction in the mesenchymal niche regulates the murine hair cycle clock. Nat Commun 11, 5114, doi:10.1038/s41467-020-18643-x (2020).

69 Hagner, A. et al. Transcriptional Profiling of the Adult Hair Follicle Mesenchyme Reveals R-spondin as a Novel Regulator of Dermal Progenitor Function. iScience 23, 101019, doi:10.1016/j.isci.2020.101019 (2020).

70 Pentinmikko, N. et al. Notum produced by Paneth cells attenuates regeneration of aged intestinal epithelium. Nature 571, 398–402, doi:10.1038/s41586-019-1383-0 (2019).

71 Brack, A. S. et al. Increased Wnt signaling during aging alters muscle stem cell fate and increases fibrosis. Science 317, 807–810, doi:10.1126/science.1144090 (2007).

72 Kusumbe, A. P. et al. Age-dependent modulation of vascular niches for haematopoietic stem cells. Nature 532, 380–384, doi:10.1038/nature17638 (2016).

73 Dulken, B. W. et al. Single-cell analysis reveals T cell infiltration in old neurogenic niches. Nature 571, 205–210, doi:10.1038/s41586-019-1362-5 (2019).

74 Ambrosi, T. H. et al. Aged skeletal stem cells generate an inflammatory degenerative niche. Nature 597, 256–262, doi:10.1038/s41586-021-03795-7 (2021).

75 Kalamakis, G. et al. Quiescence Modulates Stem Cell Maintenance and Regenerative Capacity in the Aging Brain. Cell 176, 1407–1419 e1414, doi:10.1016/j.cell.2019.01.040 (2019).

76 Yang, D. & de Haan, G. Inflammation and Aging of Hematopoietic Stem Cells in Their Niche. Cells 10, doi:10.3390/cells10081849 (2021).

77 Sugaya, M. The Role of Th17-Related Cytokines in Atopic Dermatitis. Int J Mol Sci 21, doi:10.3390/ijms21041314 (2020).

78 Miossec, P. Update on interleukin-17: a role in the pathogenesis of inflammatory arthritis and implication for clinical practice. RMD Open 3, e000284, doi:10.1136/rmdopen-2016-000284 (2017).

79 Loh, S. H., Moon, H. N., Lew, B. L. & Sim, W. Y. Role of T helper 17 cells and T regulatory cells in alopecia areata: comparison of lesion and serum cytokine between controls and patients. J Eur Acad Dermatol Venereol 32, 1028–1033, doi:10.1111/jdv.14775 (2018).

80 Ramot, Y., Marzani, B., Pinto, D., Sorbellini, E. & Rinaldi, F. IL-17 inhibition: is it the long-awaited savior for alopecia areata? Arch Dermatol Res 310, 383–390, doi:10.1007/s00403-018-1823-y (2018).

81 Sanchez-Duenas, L. E., Rojano-Fritz, L. K. & Garcia-Rodriguez, J. C. Generalized hypertrichosis associated with the use of interleukin 17 blockers in 2 patients with psoriasis. JAAD Case Rep 6, 683–685, doi:10.1016/j.jdcr.2020.05.023 (2020).

82 Yajima, M., Akeda, T., Kondo, M., Habe, K. & Yamanaka, K. Alopecia Diffusa while Using Interleukin-17 Inhibitors against Psoriasis Vulgaris. Case Rep Dermatol 11, 82–85, doi:10.1159/000499030 (2019).

83 Hao, Y. et al. Integrated analysis of multimodal single-cell data. Cell 184, 3573–3587 e3529, doi:10.1016/j.cell.2021.04.048 (2021).

84 Mereu, E. et al. Benchmarking single-cell RNA-sequencing protocols for cell atlas projects. Nat Biotechnol 38, 747–755, doi:10.1038/s41587-020-0469-4 (2020).

85 Dobin, A. et al. STAR: ultrafast universal RNA-seq aligner. Bioinformatics 29, 15–21, doi:10.1093/bioinformatics/bts635 (2013).

86 Love, M. I., Huber, W. & Anders, S. Moderated estimation of fold change and dispersion for RNA-seq data with DESeq2. Genome Biol 15, 550, doi:10.1186/s13059-014-0550-8 (2014).

87 Liao, Y., Smyth, G. K. & Shi, W. The R package Rsubread is easier, faster, cheaper and better for alignment and quantification of RNA sequencing reads. Nucleic Acids Res 47, e47, doi:10.1093/nar/gkz114 (2019).

88 Wu, D. et al. ROAST: rotation gene set tests for complex microarray experiments. Bioinformatics 26, 2176–2182, doi:10.1093/bioinformatics/btq401 (2010).

89 Xie, Z. et al. Gene Set Knowledge Discovery with Enrichr. Curr Protoc 1, e90, doi:10.1002/cpz1.90 (2021).

