## Supplementary Figures for "Local IL-17 orchestrates skin aging"

Fig. S1

a

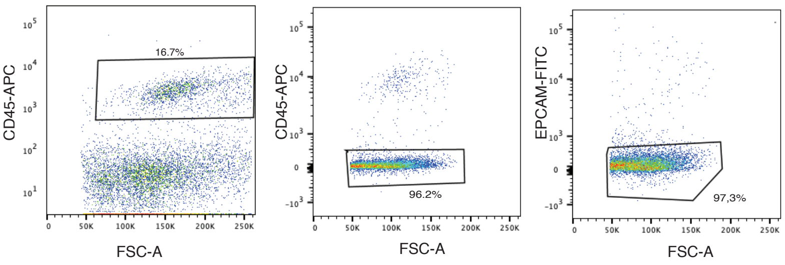

CD45-APC
FSC-A
16.7%
CD45-APC
FSC-A
96.2%
EPCAM-FITC
FSC-A
97,3%


b

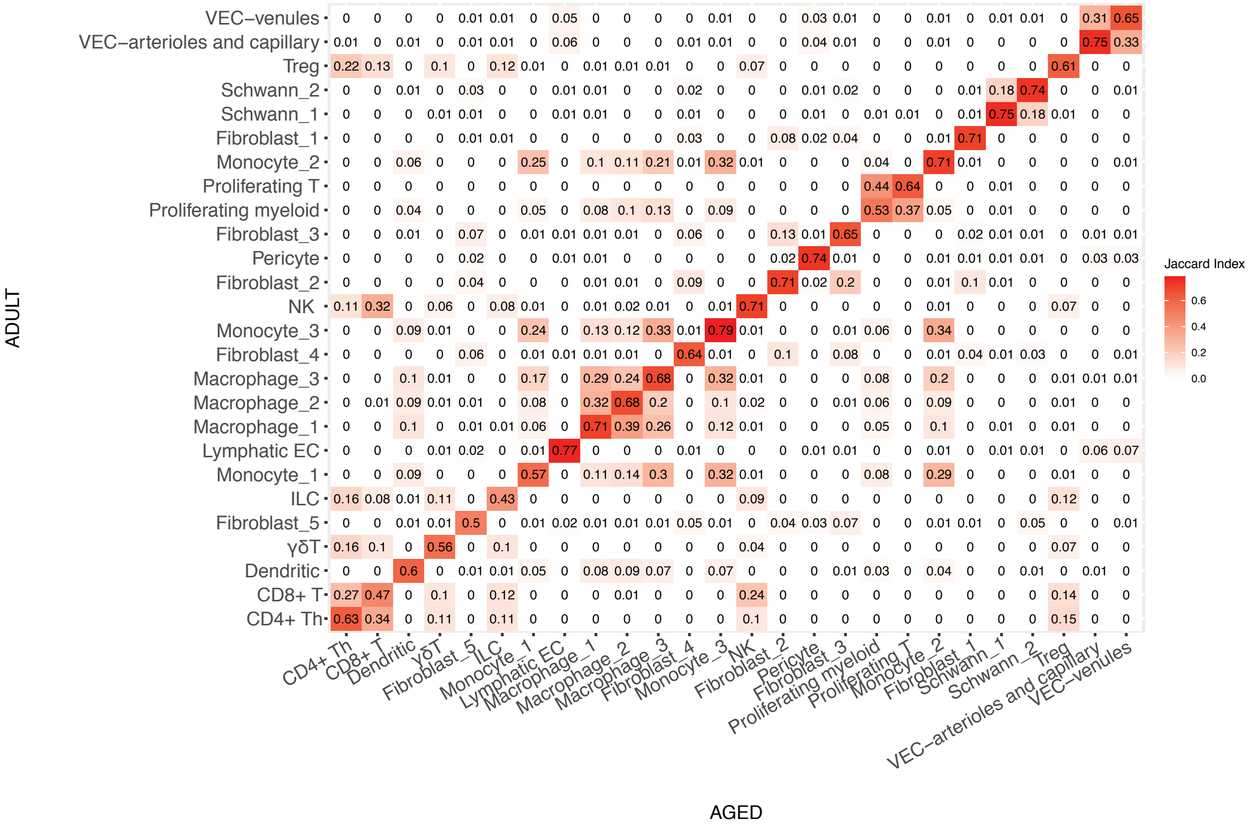

ADULT
VEC−venules
VEC−arterioles and capillary
Treg
Schwann_2
Schwann_1
Fibroblast_1
Monocyte_2
Proliferating T
Proliferating myeloid
Fibroblast_3
Pericyte
Fibroblast_2
NK
Monocyte_3
Fibroblast_4
Macrophage_3
Macrophage_2
Macrophage_1
Lymphatic EC
Monocyte_1
ILC
Fibroblast_5
γδT
Dendritic
CD8+ T
CD4+ Th
AGED
CD4+ Th
CD8+ T
Dendritic
γδT
Fibroblast_5
ILC
Monocyte_1
Lymphatic EC
Macrophage_1
Macrophage_2
Macrophage_3
Fibroblast_4
Monocyte_3
NK
Fibroblast_2
Pericyte
Fibroblast_3
Proliferating myeloid
Proliferating T
Monocyte_2
Fibroblast_1
Schwann_1
Schwann_2
Treg
VEC−arterioles and capillary
VEC−venules
Jaccard Index
0.6
0.4
0.2
0.0

Fig. S2

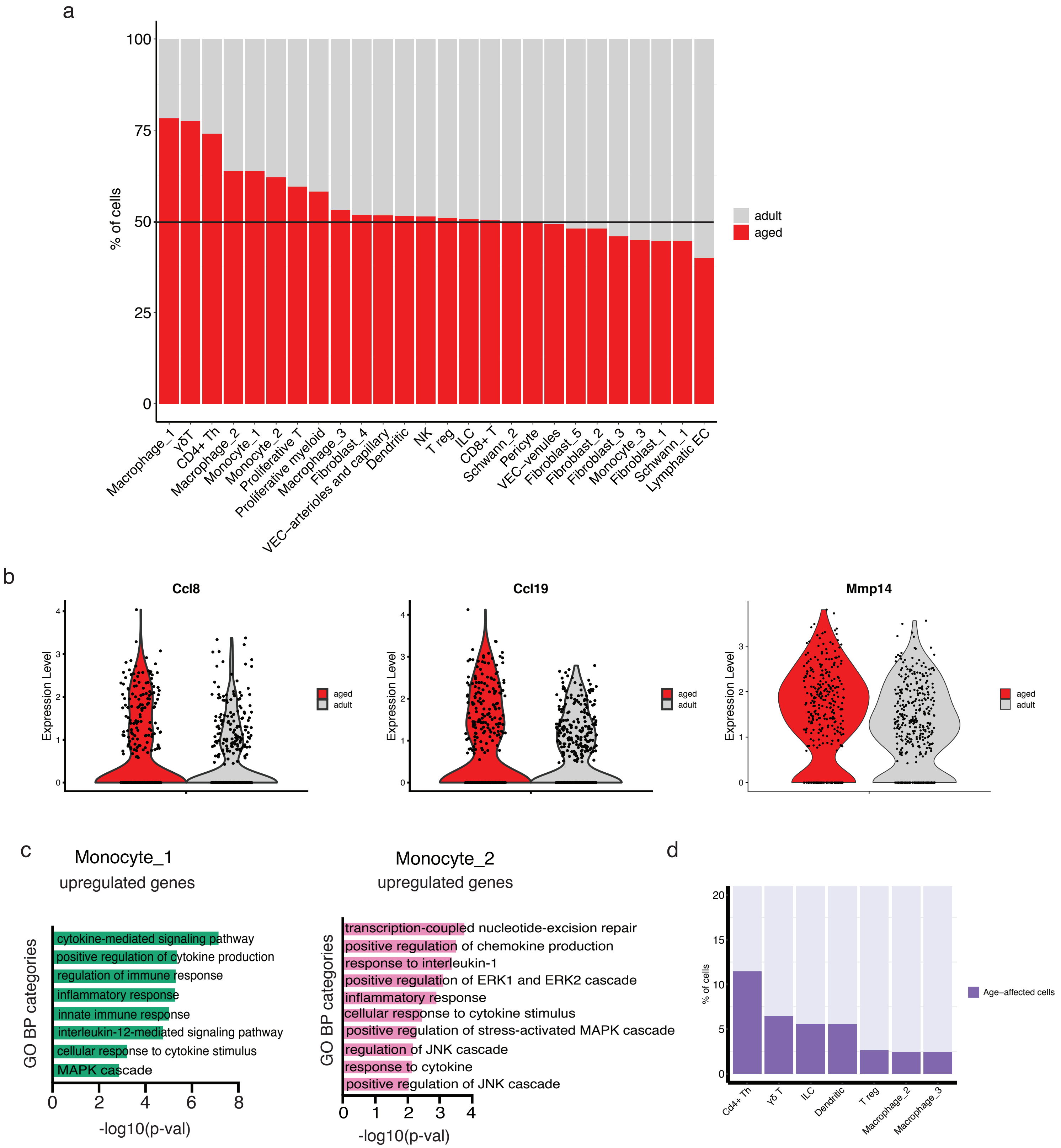
a
100
75
50
25
0
% of cells
adult
aged
Macrophage_1
γδT
CD4+ Th
Macrophage_2
Monocyte_1
Monocyte_2
Proliferative T
Proliferative myeloid
Macrophage_3
Fibroblast_4
VEC−arterioles and capillary
Dendritic
NK
T reg
ILC
CD8+ T
Schwann_2
Pericyte
VEC−venules
Fibroblast_5
Fibroblast_2
Fibroblast_3
Monocyte_3
Fibroblast_1
Schwann_1
Lymphatic EC
b
Ccl8
Ccl19
Mmp14
Expression Level
aged
adult
c
Monocyte_1
upregulated genes
GO BP categories
cytokine-mediated signaling pathway
positive regulation of cytokine production
regulation of immune response
inflammatory response
innate immune response
interleukin-12-mediated signaling pathway
cellular response to cytokine stimulus
MAPK cascade
0 2 4 6 8
-log10(p-val)
Monocyte_2
upregulated genes
transcription-coupled nucleotide-excision repair
positive regulation of chemokine production
response to interleukin-1
positive regulation of ERK1 and ERK2 cascade
inflammatory response
cellular response to cytokine stimulus
positive regulation of stress-activated MAPK cascade
regulation of JNK cascade
response to cytokine
positive regulation of JNK cascade
0 1 2 3 4
-log10(p-val)
d
20
15
10
5
0
% of cells
Age−affected cells
Cd4+ Th
γδ T
ILC
Dendritic
T reg
Macrophage_2
Macrophage_3

Fig. S3

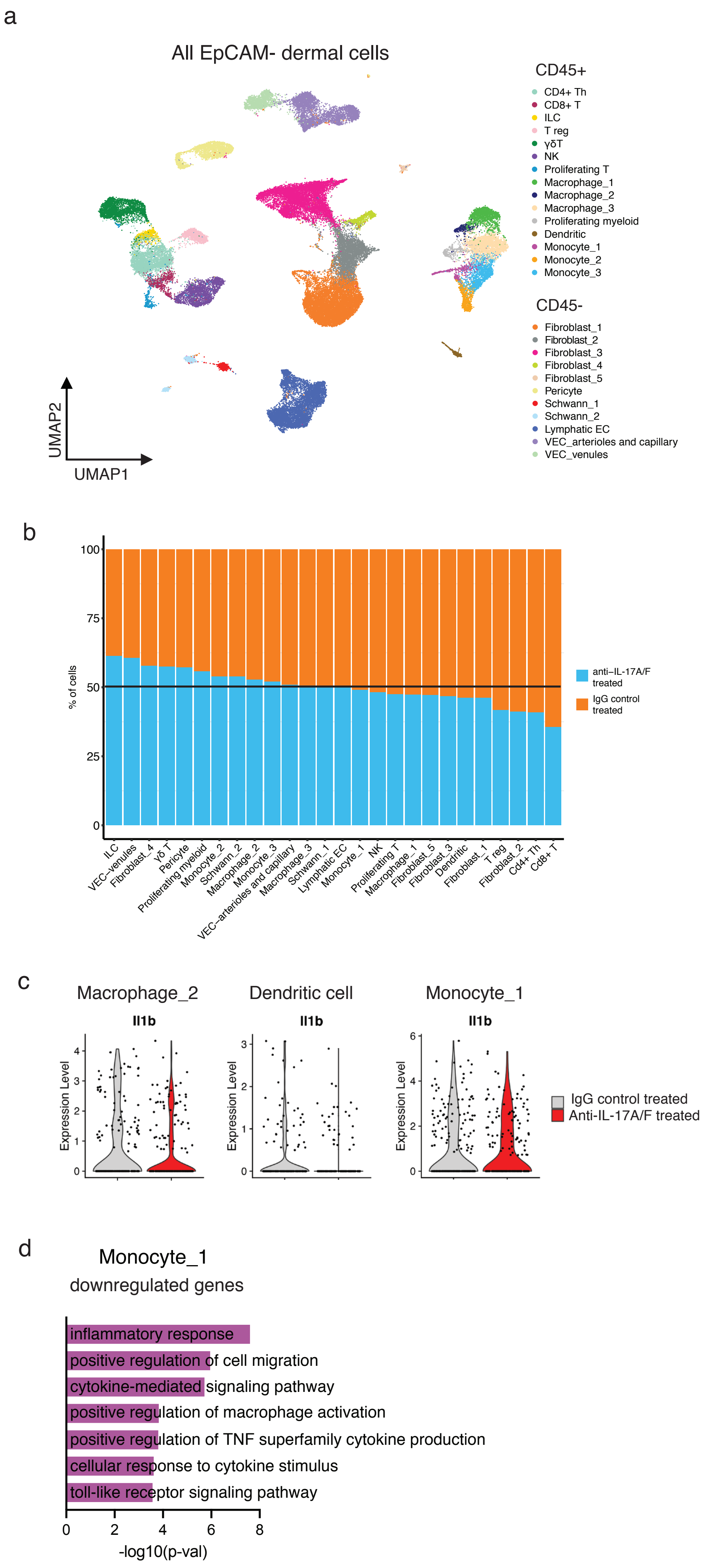
a
All EpCAM- dermal cells
CD45+
CD4+ Th
CD8+ T
ILC
T reg
γδT
NK
Proliferating T
Macrophage_1
Macrophage_2
Macrophage_3
Proliferating myeloid
Dendritic
Monocyte_1
Monocyte_2
Monocyte_3
CD45-
Fibroblast_1
Fibroblast_2
Fibroblast_3
Fibroblast_4
Fibroblast_5
Pericyte
Schwann_1
Schwann_2
Lymphatic EC
VEC_arterioles and capillary
VEC_venules
UMAP2
UMAP1
b
% of cells
100
75
50
25
0
anti−IL-17A/F treated
IgG control treated
ILC
VEC−venules
Fibroblast_4
γδ T
Pericyte
Proliferating myeloid
Monocyte_2
Schwann_2
Macrophage_2
Monocyte_3
VEC−arterioles and capillary
Macrophage_3
Schwann_1
Lymphatic EC
Monocyte_1
NK
Proliferating T
Macrophage_1
Fibroblast_5
Fibroblast_3
Dendritic
Fibroblast_1
T reg
Fibroblast_2
Cd4+ Th
Cd8+ T
c
Macrophage_2
Dendritic cell
Monocyte_1
Il1b
Expression Level
IgG control treated
Anti-IL-17A/F treated
d
Monocyte_1
downregulated genes
inflammatory response
positive regulation of cell migration
cytokine-mediated signaling pathway
positive regulation of macrophage activation
positive regulation of TNF superfamily cytokine production
cellular response to cytokine stimulus
toll-like receptor signaling pathway
0
2
4
6
8
-log10(p-val)

Fig. S4

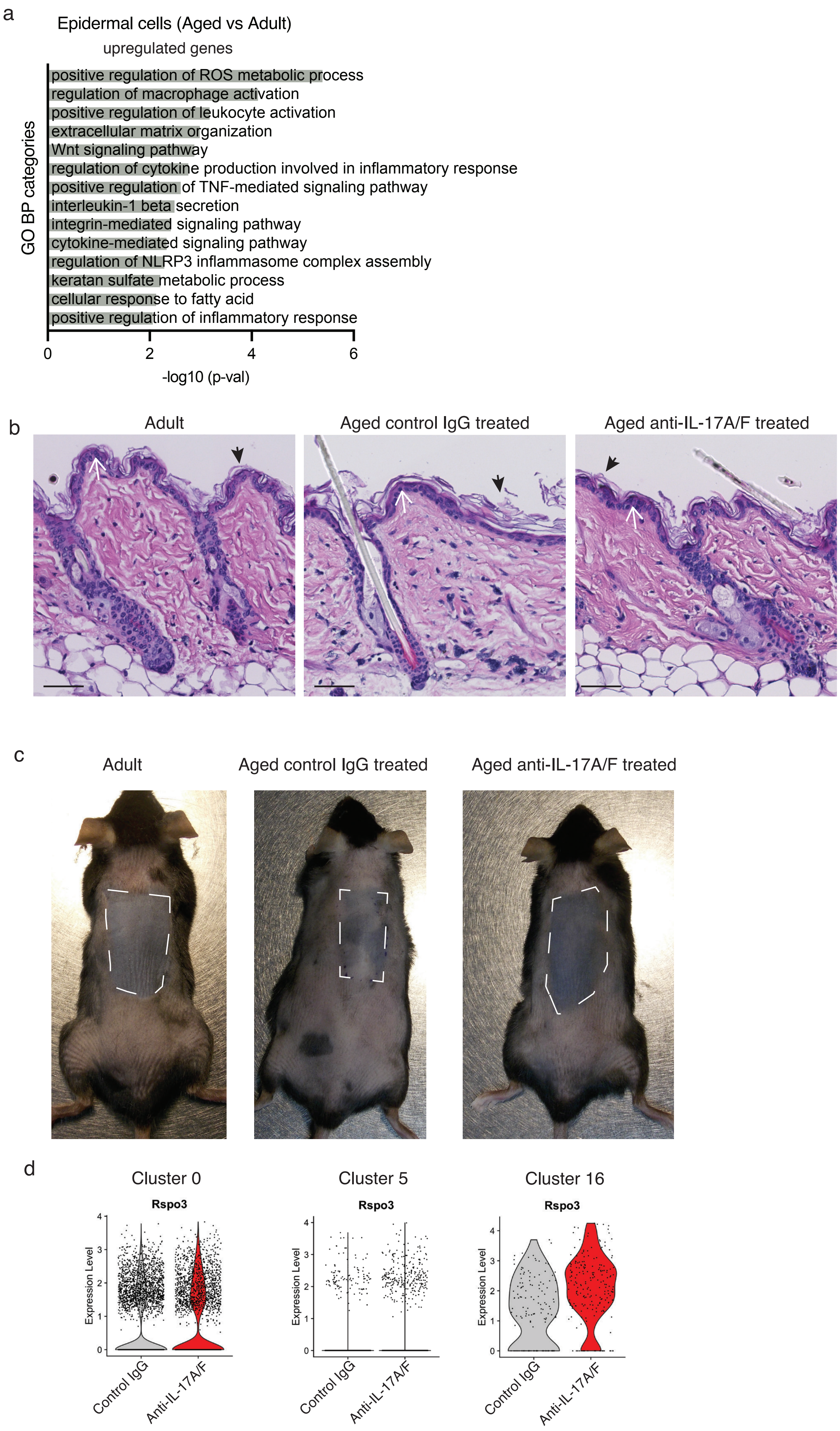
a
Epidermal cells (Aged vs Adult)
upregulated genes
GO BP categories
positive regulation of ROS metabolic process
regulation of macrophage activation
positive regulation of leukocyte activation
extracellular matrix organization
Wnt signaling pathway
regulation of cytokine production involved in inflammatory response
positive regulation of TNF-mediated signaling pathway
interleukin-1 beta secretion
integrin-mediated signaling pathway
cytokine-mediated signaling pathway
regulation of NLRP3 inflammasome complex assembly
keratan sulfate metabolic process
cellular response to fatty acid
positive regulation of inflammatory response
0
2
4
6
-log10 (p-val)
b
Adult
Aged control IgG treated
Aged anti-IL-17A/F treated
c
Adult
Aged control IgG treated
Aged anti-IL-17A/F treated
d
Cluster 0
Cluster 5
Cluster 16
Rspo3
Expression Level
Control IgG
Anti-IL-17A/F

**Supplementary Fig. 1**. **a**, Flow cytometry plots of examples for the sorting strategy to obtain CD45+ and CD45–/EpCAM– cells for 10× scRNA-seq. **b**, Plot showing the Jaccard index obtained comparing the top 100 markers of each cluster obtained in Fig. 1b between adult and aged.

**Supplementary Fig. 2**. **a,** Bar plot with the proportion of cells per cluster comparing aged and adult mice. **b,** Violin plot of Ccl8, Ccl19 and Mmp14 genes related to inflammation that are up-regulated in aged fibroblast_3 cluster. **c**, Plot of selected GO categories belonging to BP analysis for genes upregulated in monocyte_1 cluster (left panel) and monocyte_2 cluster (right panel) upon aging. The x axis represents –($\log_{10}$) of the p value for each depicted GO category. **d**, Bar plot with the percentage of cells affected by aging in each dermal cell type (only the cell types with more percentage are shown here for clarity).

**Supplementary Fig. 3**. **a**, UMAP representation of all CD45+ and CD45–/EpCAM– cells analyzed by 10× scRNA-seq after IL-17A/F blockade. **b**, Bar plot with the proportion of cells per clusters comparing IgG treated aged mice and anti-IL-17A/F treated aged mice. **c**, Violin plots showing expression values of *Il1b* in individual cells belonging to clusters macrophage_2, Dendritic cell and monocyte_1. **d**, Plot of selected GO categories belonging to BP analysis for genes downregulated in monocyte_1 upon IL-17A/F neutralization treatment. The x axis represents –($\log_{10}$) of the *P*-value for each depicted GO category.

**Supplementary Fig. 4**. **a**, Plot of selected GO categories belonging to BP analysis for genes upregulated upon aging in epidermal cells. The x axis represents –($\log_{10}$) of the *P*-value for each depicted GO category. **b**, Details of H&E staining of i) adult, ii) aged treated with control IgG and iii) aged treated with anti-IL-17A/F antibodies. Epidermal thickness (white arrows) and cornified layer thickness (black arrowheads) were quantified (Fig. 6b-e). Bar = 50 µm. **c**, Images of adult, aged/control IgG Ab and aged/anti-IL-17A/F Ab back skin showing the epilated areas (white dashed line areas). Note that the gray color of dermis indicates anagen progression of the HFs. **d,** Violin plot of *Rspo3* gene expression in cluster_0, cluster_5 and cluster_16, which subclusters correspond to the described dermal papilla fibroblasts.
